# Anterior hypothalamic parvalbumin neurons are glutamatergic and promote escape behavior

**DOI:** 10.1101/2022.09.09.507287

**Authors:** Brenton T. Laing, Megan S. Anderson, Jordi Bonaventura, Aishwarya Jayan, Sarah Sarsfield, Anjali Gajendiran, Michael Michaelides, Yeka Aponte

## Abstract

The anterior hypothalamic area (AHA) is a critical structure for defensive responding. Here, we identified a cluster of parvalbumin-expressing neurons in the AHA (AHA^PV^) that are glutamatergic with fast-spiking properties and send axonal projections to the dorsal premammillary nucleus (PMD). Using *in vivo* functional imaging, optogenetics, and behavioral assays, we determined the role of these AHA^PV^ neurons in regulating behaviors essential for survival. We observed that AHA^PV^ neuronal activity significantly increases when mice are exposed to a predator, and in a real-time place preference assay, we found that AHA^PV^ neuron photoactivation is aversive. Moreover, activation of both AHA^PV^ neurons and the AHA^PV^→PMD pathway triggers escape responding during a predator-looming test. Furthermore, escape responding is impaired after AHA^PV^ neuron ablation, and anxiety-like behavior as measured by the open field and elevated plus maze assays does not seem to be affected by AHA^PV^ neuron ablation. Finally, whole-brain metabolic mapping using positron emission tomography combined with AHA^PV^ neuron photoactivation revealed discrete activation of downstream areas involved in arousal, affective, and defensive behaviors including the amygdala and the substantia nigra. Our results indicate that AHA^PV^ neurons are a functional glutamatergic circuit element mediating defensive behaviors, expanding the identity of genetically defined neurons orchestrating fight-or-flight responses. Together, our work will serve as a foundation for understanding neuropsychiatric disorders such as aggression or fear.

## INTRODUCTION

Anterior hypothalamic area (AHA) neurons are activated by acute threat (Mitra et al., 2016) and are known to modulate panicogenic-like (Falconi-Sobrinho and Coimbra, 2018) and fighting behaviors (Ferris et al., 1997). Interestingly, the AHA is necessary for ventromedial hypothalamic nucleus (VMH)-mediated induction of defensive behavior (Fuchs et al., 1985). Whereas the AHA receives dense VMH innervation, these projections do not appear to be specific to a particular threat modality (Carvalho et al., 2020). Neurons encoding for aggression and social-fear have been identified in the mouse ventrolateral VMH (Sakurai et al., 2016) and direct activation of excitatory terminals in the AHA promotes avoidance (Wang et al., 2015), which is mediated, in part, by dorsomedial VMH steroidogenic factor 1 (NR5A1 or SF1)-expressing neurons (Wang *et al*., 2015). Thus, these studies indicate that the AHA may play a key role in regulating the arousal or avoidance associated with defensive behaviors. However, the specific neuronal types within the AHA encoding for such behaviors have yet to be determined.

Within the mouse AHA, a cluster of parvalbumin-expressing neurons (PVALB; AHA^PV^) has been implicated in the central autonomic control of blood pressure and heart rate, particularly in response to thermal stressors to modulate autonomic arousal (Mittag et al., 2013). Therefore, we sought to determine whether AHA^PV^ neurons represent a discrete neuronal subtype involved in regulating defensive behaviors.

We first used *in vivo* functional imaging to determine whether AHA^PV^ neurons are active during a predator exposure test and followed this with the use of machine learning models to predict the animals’ behavioral states. Next, we characterized the electrophysiological properties and expression of specific ion channels of AHA^PV^ neurons using brain slice electrophysiology and *in situ* hybridization techniques. Moreover, neuronal tracing methods and Channelrhodopsin-2-assisted circuit mapping (CRACM) were used to determine AHA^PV^ axonal projections. Subsequently, we manipulated the activity of AHA^PV^ neurons and the AHA^PV^→PMD pathway by using optogenetics during reward- and threat-related behavioral assays, and we specifically ablated AHA^PV^ neurons to examine whether their activity is necessary for escape responding and anxiety-like behavior. Finally, we performed whole-brain metabolic mapping to identify brain regions activated during the stimulation of AHA^PV^ neurons by combining positron emission tomography (PET) and optogenetics.

## MATERIALS AND METHODS

### Animals

All experimental protocols were conducted in accordance with U.S. National Institutes of Health Guidelines for the Care and Use of Laboratory Animals and with the approval of the National Institute on Drug Abuse Care and Use Committee. Male and female heterozygous *Pvalb^Cre^*mice (C57BL/6J background; IMSR Cat# JAX:008069, RRID:IMSR_JAX:008069; The Jackson Laboratory, ME, U.S.A.) and *Slc32a1^Cre^* mice (C57BL/6J background; IMSR Cat# JAX:028862, RRID:IMSR_JAX:028862; The Jackson Laboratory) were used. For characterization of the electrophysiological properties of AHA^PV^ neurons, the *Pvalb^Cre^* line was crossed with *Rosa26^LSL-tdtomato^* (C57BL6/J background; IMSR Cat# JAX:007909, RRID:IMSR_JAX:007909) to generate *Pvalb^Cre/+^;Rosa26^LSL-tdtomato/LSL-tdTomato^*mice. For *in situ* hybridization experiments, wildtype mice (C57BL/6J background; ISMR Cat# JAX:000644, RRID:ISMR_JAX:000664) were used. For *in vivo* functional imaging during threat response, adult male Wistar rats (RRID:RGD_13508588; Strain # 003, Charles River Laboratories, MA, U.S.A.) were used as the predator threat. Rodents were group housed with littermates in temperature and humidity-controlled rooms on a 12 h light/dark cycle with *ad libitum* access to water and rodent chow (PicoLab Rodent Diet 20, 5053 tablet, LabDiet/Land O’Lakes Inc., MO, U.S.A.).

### Stereotaxic viral injections

Stereotaxic microinjections were conducted as previously described (Atasoy et al., 2008; Siemian et al., 2019). For experiments targeting the anterior hypothalamic area (AHA) an adeno-associated virus (AAV; 2 × 12.5 nL for optogenetic experiments; 5 nL for anterograde tracer cohort 1; 25 nL for anterograde tracer cohort 2) was injected into the AHA (AP: −0.70, ML: ±0.425, DV: −5.5 and −5.3 for optogenetic experiments, −5.3 for anterograde tracer experiments). Optical fibers (200-µm core, 0.48 NA, 4.8-mm length) were inserted at 5-degree angles (AP: −0.85, ML: ±0.84, DV: −4.8) dorsal to the AHA. For retrograde tracing experiments, 25 nL of 4% Fluoro-Gold (Fluorochrome LLC, Denver, CO) were injected into the dorsal premammillary nucleus (AP: −2.55, ML: +0.70, DV: −5.5). As a surgical control for caspase experiments, control mice were injected with a Cre-dependent AAV expressing the fluorophore tdTomato. For *in vivo* functional imaging experiments, a GRIN lens (500 µm, Snap-in imaging cannula Model L-V; Doric Lenses, Inc., Québec, QC, Canada) was implanted to record AHA neuronal activity (AP: −0.85, ML: +0.425, DV: −5.3) as described previously (Siemian et al., 2021).

Viruses used include:

(1) rAAV2/9-CAG-FLEX-jGCaMP8f-WPRE, titer: 5.0 × 10^12^ GC/ml (Addgene viral prep # 162382-AAV9, RRID:Addgene_162382; Addgene, MA, U.S.A.),
(2) rAAV2/9-EF1a-double floxed-hChR2(H134R)-EYFP-WPRE-HGHpA, titer: 5.0 × 10^12^ GC/ml (Addgene viral prep # 20298-AAV9, RRID:Addgene_20298),
(3) rAAV2/9-EF1a-DIO-YFP, titer: 5.0 × 10^12^ GC/ml (Addgene viral prep # 27056-AAV9, RRID:Addgene_27056),
(4) rAAV2/8-EF1a-DIO-iC++-EYFP, titer: 5.0 × 10^12^ GC/ml (Cat # GVVC-AAV-108; Stanford Neuroscience Gene Vector and Virus Core, CA, U.S.A.),
(5) rAAV2/1-EF1a-FLEX-taCasp3-TEVp, titer: 1.9 × 10^12^ GC/ml (RRID:Addgene_45580; University of North Carolina Vector Core NC, U.S.A.),
(6) rAAV2/1-CAG-FLEX-EGFP-WPRE, titer: 5.0 × 10^12^ GC/ml (Cat # AV-1-ALL854, RRID:Addgene_51502; University of Pennsylvania (U Penn) Vector Core, PA, U.S.A.),
(7) rAAV2/1-EF1a-DIO-synaptophysin (SYP)-mCherry, titer: 5.0 × 10^12^ GC/ml (Cat # AAV-RN1; Massachusetts General Hospital Gene Delivery Technology Core, MA, U.S.A.),
(8) rAAV2/1-CAG-FLEX-ChR2-tdTomato, titer: 5 × 10^12^ GC/ml (Addgene viral prep # 18917-AAV1, RRID:Addgene_18917), and
(9) rAAV2/1-CAG-FLEX-tdTomato, titer: 4.5 × 10^13^ GC/ml (Cat # AV-1-ALL864, RRID:Addgene_51503; U Penn Vector Core).

### *In vivo* functional imaging and predatory threat exposure

For *in vivo* functional imaging the Cre-dependent genetically encoded calcium indicator jGCaMP8f was used for visualization of neuronal activity in *Slc32a1^Cre^* and *Pvalb^Cre^* mice. Mice were habituated to experimenter handling, behavioral rooms, and miniscope tethering for up to 1 h per day for 3 days prior to testing, and tests were conducted within the light phase of the light/dark cycle. The Doric Lenses, Inc. Basic Fluorescence Snap-in Microscopy System and Doric Neuroscience Studio software v5.1 (RRID:SCR_018569) were used for image acquisition.

A predatory threat exposure paradigm (Esteban Masferrer et al., 2020) was adapted for *in vivo* functional imaging experiments. Adult male Wistar rats were used as predators. The arena consisted of an opaque black, H-shaped box with bedding on the floor; a transparent wall with holes for olfaction was placed in the hallway to enclose one portion of the box, keeping the rat and mouse physically separated from each other.

The test consisted of 4 × 3 min epochs (baseline, threat exposure, assessment, and exploration), and the mouse was briefly removed and placed in an intermediate holding cage between all epochs. First, the mouse was placed into the apparatus and a baseline was acquired while the rat remained in a separate location. Second, the threat exposure stage was conducted by placing a rat in the enclosed side of the arena followed by placement of the mouse on the side where the baseline was conducted. For the third stage (assessment), the rat was removed from the room, and the mouse was re-introduced to the original/baseline side. Finally, for the exploration stage, the transparent wall was removed and the mouse was re-introduced with access to explore the predator side of the apparatus.

### Brain slice preparation and electrophysiology

Mice were deeply anesthetized with isoflurane and then transcardially perfused with an *N*-methyl-D-glucamine solution containing (in mM): 92 NMDG, 20 HEPES, 25 D-glucose, 30 NaHCO_3_, 1.2 NaH_2_PO_4_, 2.5 KCl,5 sodium ascorbate, 3 sodium pyruvate, 2 thiourea, 10 MgSO_4_, and 0.5 CaCl_2_. Prior to carbogenation with 95% O_2_ and 5% CO_2_, all external solutions were adjusted to pH 7.4 and 305 − 310 mOsm. After visualization of perfusion-induced color-change in the liver, mice were decapitated, and brains were removed. 200-µm coronal sections were obtained using a Leica VT1200 S vibratome (RRID:SCR_020243). Brain slices were transferred to a warm bath (32 °C) until approximately 12-min post-decapitation, and then, slices were transferred to a Brain Slice Keeper 4 (Cat # S-BSK4; AutoMate Scientific, CA, U.S.A.) filled with room temperature HEPES-based holding solution containing (in mM): 92 NaCl, 20 HEPES, 25 D-glucose, 30 NaHCO_3_, 1.2 NaH_2_PO_4_, 2.5KCl, 5 sodium ascorbate, 3 sodium pyruvate, 2 thiourea, 1 MgSO_4_, and 2 CaCl_2_. For electrophysiological recordings to characterize the properties of AHA^PV^ neurons, a single slice was submerged in 32 °C aCSF containing (in mM): 125 NaCl, 2.5 KCl, 1.25 NaH_2_PO_4_, 1 MgCl_2_6H_2_O, 11 D-glucose, 26 NaHCO_3_, 2.4 CaCl_2_. aCSF was continuously delivered to the recording chamber by a peristaltic pump (World Precision Instruments, FL, U.S.A.) at a flow rate of 1−2 ml/min.

Characterization of the electrophysiological properties of AHA^PV^ neurons (*n* = 30) was conducted using *Pvalb^Cre/+^;Rosa26^LSLtdTomato/LSLtdTomato^*mice. Neurons were identified using epifluorescence and differential interference contrast (IR-DIC) optics with an AxioExaminer.Z.1 microscopy (Carl Zeiss Microscopy LLC). pCLAMP v10.3 software (RRID:SCR_011323; Molecular Devices LLC, CA, U.S.A.) was used for acquisition with a Multiclamp 700B amplifier and a Digidata 1440A (10-kHz digitization for current clamp recordings, 50 kHz for voltage clamp). Borosilicate glass patch pipettes (3.5−4.5 MΩ) were pulled using a P-2000 laser-based micropipette puller (Sutter Instruments, CA, U.S.A.). Current clamp recordings were conducted using a potassium gluconate based internal solution (in mM): 135 potassium gluconate, 10 HEPES, 4 KCl, 4 MgATP, 0.3 NA_3_GTP. The potassium gluconate solution was adjusted to pH 7.3 (using KOH) and 290 mOsm. Gigaohm seals were obtained using a holding potential of −70 mV prior to breaking into the cell. For voltage clamp recordings, a −2 mV hyperpolarizing injection was delivered on each sweep to monitor access to the cell, and only stable recordings were included in analysis. Electrophysiological analysis of intrinsic membrane properties and spike characteristics were conducted as previously described (Kisner et al., 2018). Channelrhodopsin-2-assisted circuit mapping (CRACM) experiments were conducted by obtaining whole cell access and monitoring post-synaptic currents in voltage clamp using a CsMs based internal solution (in mM: 117 cesium methanesulfonate, 20 HEPES, 0.4 EGTA, 2.8 NaCl, 5 TEA-Cl, 4 Mg-ATP, 0.4 Na-GTP; pH adjusted to 7.3 using CsOH; 289 mOsm) with excitatory post-synaptic currents (EPSCs) checked at −70 mV. For a subset of cells, cyanquixaline (CNQX, 10 µM) was bath applied to block AMPA receptor-mediated currents. For a subset of cells not subjected to any pharmacological manipulation, inhibitory post-synaptic currents (IPSCs) were checked with a holding potential at 0 mV.

### Fluorescent *in situ* hybridization

After cervical dislocation, brains from wildtype mice were dissected and rapidly frozen in −80 °C isopentane, then subsequently stored at −80 °C. Coronal cryosections (20 µm) containing the AHA (bregma −0.45 to −1.35) were sliced using a Leica CM3050 S cryostat (Leica Biosystems Inc.) and sections were collected onto Superfrost Plus glass slides (VWR International). Slides were stored at −80 °C prior to processing. Fluorescent *in situ* hybridization was performed using the RNAscope® Multiplex Fluorescent Assay v1 for fresh frozen tissue (Cat # 320851; Advanced Cell Diagnostics Inc., CA, U.S.A.). Briefly, sections were fixed in 4% PFA in PBS, dehydrated by ethanol series, and treated with Protease IV. Sections were incubated with target probes for mouse *Pvalb* (Cat # 421931; accession number NM_013645.3, target region aa2–885), *Hcn2* (Cat # 427001-C2; accession number NM_008226.2, target region aa687–1878), and *Kcnc1* (Cat # 429181-C3; accession number NM_008421.3, target region aa2957–3952). After hybridization, a series of signal amplification steps (Amp1, Amp2, and Amp3) were performed per kit protocol followed by incubation with labels (Amp4C) for fluorescent visualization of each probe: *Pvalb* (Atto550), *Hcn2* (Atto647), and *Kcnc1* (Alexa488). Slides were counterstained with DAPI and coverslipped with Fluoromount-G aqueous mounting medium (Cat # 17984-25; Electron Microscopy Systems). Images were acquired using a Keyence BZ-X710 microscope (Keyence Corp., IL, U.S.A.).

Cell counts were performed on every sixth brain slice, and AHA neurons were assessed for co-expression of *Pvalb* with *Hcn2* and *Kcnc1* using a custom ImageJ macro series called Autocount (Laing, 2022; Schindelin et al., 2012). This macro series identifies all DAPI-positive areas as individual regions of interest and measures fluorescence intensity across each channel. Thresholds were set for each channel to be designated as positive or negative for each marker.

### Radioactive *in situ* hybridization and immunohistochemistry

Wildtype mice were deeply anesthetized with isoflurane and transcardially perfused with 0.1 M phosphate buffer (PB) followed by 4% PFA in 0.1 M PB. Whole brains were removed and post-fixed in 4% PFA/PB for 2 h at 4 °C. Samples were washed with 0.1 M PB (2 × 30 min each) at 4 °C, and then transferred to 18% sucrose in 0.1 M PB. Free-floating coronal cryosections (14 µm) were sliced using a Leica CM3050 S cryostat. *In situ* hybridization was performed as previously described (Kisner *et al*., 2018; Qi et al., 2016; Root et al., 2014). Steps are at room temperature unless otherwise stated. Sections were incubated for 3 × 10 min in 0.1 M PB/0.5% Triton X-100, rinsed with 0.1 M PB (3 × 10 min), treated with 0.2 N HCl for 15 min, rinsed with 0.1 M PB (3 × 10 min), and then acetylated in 0.25% acetic anhydride/0.1 M triethanolamine pH 8.0 for 10 min. Sections were rinsed for 3 × 10 min with 0.1 M PB, fixed with 4% PFA/PB for 10 min, washed again with 0.1 M PB (3 × 10 min), and then prehybridized for 2 h at 55 °C. Sections were hybridized for 16 h at 55 °C with [^35^S]- and [^33^P]-labeled (10^7^ c.p.m./ml) antisense probe for either *Vglut2* (nucleotides 317-2357; Accession Number NM_053427) or *Vgat* (nucleotides 1-2814, Accession Number BC052020). After hybridization, sections were incubated in 2× SSC buffer/10 mM β-mercaptoethanol (BME) for 30 min at room temperature. Sections were next treated with 5 µg/ml RNase A in 10 mM Tris-HCl pH 7.9/10 mM NaCl/0.1 mM EDTA for 1 h at 37 °C, washed in 0.5× SSC/50% formamide/10 mM BME/0.5% sarkosyl for 2 h at 55 °C, washed in 0.1× SSC/10 mM BME/0.5% sarkosyl for 1 h at 60 °C, and rinsed with 0.1 M PB (3× 10 min) prior to parvalbumin immunolabeling.

For immunohistochemistry, sections were blocked with 0.1 M PB/4% bovine serum albumin (BSA)/3% Triton X-100 for 1 h. Sections were incubated with goat anti-parvalbumin antibody (1:1000 PVG-213, Swant) in block solution overnight at 4 °C. After washing with 0.1 M PB (3 × 10 min), sections were incubated with biotinylated anti-goat IgG secondary antibody (1:200) in block solution, washed with 0.1 M PB (3 × 10 min), incubated for 1 h in avidin-biotinylated horseradish peroxidase (1:200, ABC kit; Vector Laboratories, CA, U.S.A.), rinsed with 0.1 M PB (3 × 10 min), and developed with 0.05% 3,3-diaminobenzidine-4 HCl (DAB)/0.003% hydrogen peroxide/0.1 M PB for 10 min. Sections were washed with 0.1 M PB (3 × 10 min), mounted onto coated slides, dipped in Ilford K5 nuclear tract emulsion (Harman Technology Ltd, TX, U.S.A.), and exposed in the dark at 4 °C for four weeks prior to development.

Sections were imaged with brightfield and epiluminescence microscopy using a Leica DMR microscope with a 20× objective and cellSens Standard v1.11 software (Olympus Corporation). Neurons observed within the AHA^PV^ region were manually assessed for the co-expression of *Vglut2* mRNA or *Vgat* mRNA with anti-parvalbumin immunolabeled cells. Brightfield was used to determine whether a parvalbumin-immunolabeled (brown DAB product) neuron contained the aggregates of silver grains for *Vglut2* mRNA or *Vgat* mRNA, which were viewed under epiluminescence.

### Optogenetic manipulations

For testing, mice were tethered to a 450-nm laser (Doric Lenses, Inc.) via optical fiber cables. Light stimulation and inhibition protocols were generated using Doric Neuroscience Studio software v5.1 using the following parameters: (1) ChR2/YFP photoactivation, 10 − 15 mW illumination for 10-ms pulse width at 20 Hz and (2) iC++/YFP photoinhibition, 9-s pulse width at 0.1 Hz.

### Behavioral assays

All behavioral tests were conducted within the light phase of the light/dark cycle. Mice were acclimated to the testing room for at least 1 h prior to each experiment. Videos were recorded within ANY-maze video tracking software v7 (RRID:SCR_014289; Stoelting Co., IL, U.S.A.) unless otherwise noted.

#### Real-time place preference (RTPP)

For real-time place preference experiments, a standard-sized rat cage (20 cm × 40 cm) with black opaque walls and a layer of bedding was placed in an isolation chamber with the overhead lights turned off. Photoactivation or photoinhibition was paired with one side of the chamber (laser-ON side) and was consistent across sessions. Mice were placed in the laser-OFF side and could freely transition between the two sides for 20 minutes. Average speed and total time spent in each side of the chamber were calculated by ANY-maze software. Immobility was defined as the absence of movement in the X, Y, and Z space (Wang *et al*., 2015). For immobility detection, the sensitivity slider was set to 80% with a minimum duration threshold of 2 seconds.

#### Predator-looming test

A simulated predator threat exposure assay was conducted as previously described (Yilmaz and Meister, 2013). Briefly, a 24-inch digital screen was placed above an open apparatus (40 cm × 40 cm). The looming stimulus was a black circle on a gray background that increased in size and then disappeared. A safety zone (nest) consisting of a tent was placed in one corner of the arena, and the start zone was set in the opposite corner. For optogenetic experiments, photoactivation or photoinhibition began simultaneously with the predator-looming stimulus, and mice were provided an additional 2 days (10 min/day) of habituation to the apparatus while tethered to fiber optic patch cords. This was critical for facilitation of claiming the nest, which was less natural than in the untethered mice used for the genetic ablation experiments.

#### Open field test

For the open field test, mice were placed in open field arenas for 18 min. Each arena (30 cm × 30 cm) had a thin layer of bedding and was inside an isolation chamber illuminated to approximately 150 lx. An overhead camera and ANY-maze software were used to record and assess locomotion and location.

#### Light-dark transition test

For light-dark transition testing, a square testing arena (30 cm × 30 cm) was divided into two zones (‘dark zone’: 10 cm × 30 cm and ‘light zone’: 20 cm × 30 cm) using an opaque black insert that had a small hole for the mouse to pass through. The arena was placed into an isolation chamber, and the light zone was illuminated to approximately 400 lx. Mice could freely transition between the light and dark zones for 5 minutes. ANY-maze software was used to track the number of zone transitions and the total time spent in each zone.

#### Elevated plus maze

The elevated plus maze was a standard apparatus consisting of two open arms (30 × 5 cm) and two closed arms (30 × 5 × 30 cm) extending from a central platform (5 × 5 cm). The apparatus was elevated 75 cm from the floor. The room was illuminated to 300 lx via overhead lighting. Mice were placed onto the center platform and allowed to freely explore the maze for 5 min. EthoVision XT v8.5 software (RRID:SCR_000441; Noldus, Wageningen, Netherlands) was used to record and track mice.

### Positron Emission Tomography (PET)

Mice were injected with AAV-FLEX-ChR2-tdTomato or AAV-FLEX-tdTomato into the AHA and unilateral optical fibers were implanted dorsal to the AHA in the right hemisphere. Mice were fasted for 16 h before the experiment. On the day of the experiment, mice were anesthetized with 1.5% isoflurane and placed on a custom-made bed in a nanoScan small animal PET/CT scanner (Mediso Medical Imaging Systems, Budapest, Hungary). Mice were then injected (i.p.) with 13 MBq of 2-deoxy-2-[18F]fluoro-D-glucose (FDG; Cardinal Health, Inc., OH, U.S.A.) and scanned for 30 min using a dynamic acquisition protocol followed by a CT scan. During FDG uptake mice were photostimulated using 3-min OFF/ON blocks (10−15 mW, 10-ms pulse, 20 Hz). PET data were reconstructed and corrected for dead-time and radioactive decay (Thanos et al., 2013b). All qualitative and quantitative assessments of PET images were performed using the PMOD software environment (RRID:SCR_016547; PMOD Technologies LLC, Zurich, Switzerland) and Mediso’s Nucline software. The data were reconstructed in frames corresponding to the blocks of stimulation and the dynamic PET images were co-registered to magnetic resonance imaging (MRI) templates using PMOD’s built-in atlases. All statistical parametric mapping analyses were performed using MATLAB (R2016a; RRID:SCR_001622) and SPM12 (RRID:SCR_007037; https://www.fil.ion.ucl.ac.uk/spm/software/spm12/; University College London, London, U.K.).

### Histology

Mice were deeply anesthetized with isoflurane and transcardially perfused with 1x phosphate buffered saline (PBS) followed by 4% paraformaldehyde (PFA) in 1x PBS. Whole brains were removed and were post-fixed in 4% PFA at 4°C for at least 24 h before further processing. For optogenetic and *in vivo* imaging experiments, tissue was embedded in 4% agarose in PBS and 50-µm free-floating, coronal brain sections were collected using a vibratome (Leica VT1200S vibrating microtome, RRID:SCR_020243; Leica Biosystems Inc., IL, U.S.A.). For neuronal ablation experiments, samples were cryoprotected in 30% sucrose in 1x PBS, frozen on dry ice, and mounted in Cryo-Gel Tissue Embedding Medium (Leica Biosystems GmBH, Wetzlar, Germany). Coronal brain sections (20-µm thick) were collected in 1x PBS using a Leica Biosystems CM3050 S cryostat. For all experiments, sections were mounted with DAPI-Fluoromount-G aqueous mounting medium (Cat # 17984-24, Electron Microscopy Sciences, PA, U.S.A.) onto Superfrost Plus glass slides (Cat # 48311-703, VWR International, PA, U.S.A.) and imaged with an AxioZoom.V16 fluorescence microscope (Carl Zeiss Microscopy, NY, U.S.A.).

For optogenetic experiments and anterograde tracer experiments, the AHA was imaged, and mice were excluded if there was no virus expression in the AHA. If fibers were implanted over the PMD, slices between bregma −2.3 to −2.6 were imaged to visualize axons and optical fiber placement in the PMD. For Fluoro-Gold experiments, mice were excluded if Fluoro-Gold expression was not localized to the PMD.

For verification of genetic ablation of AHA^PV^ neurons, the absence of parvalbumin-positive cells in the AHA was assessed by immunostaining with guinea pig anti-PVALB polyclonal antibody (1:1000, Cat # GP72, RRID:AB_2665495; Swant, Marly, Switzerland). Every fifth slice between bregma −0.5 and −1.5 was imaged and manually counted. We set inclusion criteria for the AHA^PV^:Casp3 group at two-S.E.M. below the mean of the control group. Three of eleven AHA^PV^:Casp3 mice failed to meet the criteria and were excluded from subsequent behavior analyses (*n* = 8/11 included); one control AHA^PV^:mCherry animal was determined to be below this threshold and was also excluded from analysis (*n* = 7/8 included).

Immunostaining was conducted for axonal projections and verification of virus expression using a chicken anti-GFP polyclonal antibody (1:1000, Cat # GFP-1020, RRID:AB_10000240; Aves Labs, CA, U.S.A.) or a rabbit anti-DsRed polyclonal antibody (1:1000, Cat # 632496, RRID:AB_10013483, Takara Bio U.S.A., Inc., CA, U.S.A.). In addition, FOS immunostaining was conducted using rabbit anti-phospho-FOS monoclonal antibody (1:1000, Cat # 5348, RRID:AB_10557109; Cell Signaling Technology, MA, U.S.A.). Slices were washed in PBS 6 × 10 min each. Then, slices were blocked for 1 h in PBS + 0.3% Triton X-100 + 3% normal donkey serum. Samples were incubated overnight at room temperature in primary antibody diluted in block solution. The next day, slices were washed 6 × 10 min each followed by incubation in secondary antibody in block solution: goat anti-chicken Alexa Fluor 488 (1:500, Cat # A11039, RRID:AB_2534096; Thermo Fisher Scientific, MA, U.S.A.) or donkey anti-rabbit Alexa Fluor 647 (1:500, Thermo Fisher Cat # A31573, RRID:AB_2536183). For all samples, sections were mounted with DAPI-Fluoromount-G aqueous mounting medium (Electron Microscopy Sciences) onto Superfrost Plus glass slides (VWR International). All wide-field images were taken with an AxioZoom.V16 fluorescence microscope (Carl Zeiss Microscopy). Images for co-localization of AHA^PV^ neurons and retrogradely traced Fluoro-Gold were acquired using an LSM700 laser scanning confocal microscope (Carl Zeiss Microscopy).

### Experimental design and statistical analysis

Data are reported as mean ± s.e.m unless otherwise noted. Data from electrophysiological experiments were computed using Clampfit v10.6 (pCLAMP, RRID:SCR_011323; Molecular Devices LLC, CA, U.S.A.). Calcium imaging signals were extracted and analyzed using CNMF-E (Zhou et al., 2018). To predict mouse behavioral stage based on AHA^PV^ neuronal activity during the predatory threat exposure assay, the MATLAB machine learning toolbox (R2020a, RRID:SCR_001622; MathWorks, MA, U.S.A.) was trained on 80% of the cells to generate cubic KNN, cosine KNN, and quadratic SVM models to predict mouse behavioral stage based on AHA^PV^ and AHA^VGAT^ neuronal activity during exposure to a predatory threat.

GraphPad Prism 8 software (RRID:SCR_002798; GraphPad, La Jolla, CA, U.S.A.) was used for graphs and statistical analyses. Student’s *t*-test was used to determine differences between groups. For Student’s *t*-tests, Welch’s correction was applied when variance was different between two groups. Two-way repeated measures ANOVA was used to detect differences between groups and within groups when appropriate. Sphericity was not assumed and Greenhouse-Geiser corrections were made for all experiments. Sidak’s multiple comparisons tests were used for further evaluation when significant main effects were detected. The number of experimental units is depicted as ‘*n*’ for each experiment.

For experimental design, each behavioral experiment was done with an independent cohort of mice. Littermate controls were assigned to either an experimental group (ChR2, ChR2→PMD, iC++, or Casp3) or a control group (YFP or mCherry). Mice underwent stereotaxic surgery as described above and were allowed to recover for at least two weeks for optogenetic experiments and three weeks for genetic ablation experiments. For tracing experiments, each mouse is considered to be one ‘*n*’. For electrophysiological characterization and functional imaging, each individual cell was considered an experimental unit. For CRACM experiments, because photostimulation is temporally confined, each individual cell is considered an experimental unit. For PET analysis, each individual mouse is considered an experimental unit for calculation of between-group differences.

## RESULTS

### Exposure to a predatory threat triggers an increase in AHA^PV^ neuronal activity

The AHA is mainly comprised of GABAergic neurons (Boudaba et al., 1996). However, previous studies have supported the idea that different cell types in the AHA encode distinct aspects of defensive responses to threat (Bang et al., 2022; Wang *et al*., 2015). Therefore, we first sought to determine whether AHA^PV^ neuronal activity is modulated by exposure to a predatory threat such as a rat. For this, we injected a Cre-dependent adeno-associated virus (AAV) expressing the genetically encoded calcium indicator jGCaMP8f ((Zhang, 2020) into the AHA of *Slc32a1^Cre^* and *Pvalb^Cre^* mice to compare changes in neuronal activity during a predatory threat exposure paradigm (Esteban Masferrer *et al*., 2020) in AHA^VGAT^ and AHA^PV^ neurons, respectively (**Figure 1A–D**). Mice were subjected to 4 x 3 min epochs to image neuronal activity during the following stages: (1) baseline, (2) threat exposure (i.e., rat exposure), (3) post-threat assessment, and (4) exploration of the threat-related space. Signals were extracted and processed for Z-score calculation across experimental phases (Friedrich et al., 2017; Pnevmatikakis and Giovannucci, 2017). For AHA^VGAT^ neurons (*n* = 83 neurons from 3 mice), we did not detect any significant differences of Z-scored activity levels between stages (**Figure 1E–G**). Notably, a quadratic support vector machine (SVM) model trained on 80% of the cells was able to predict the behavioral stage with accuracy well above chance (56.8% compared to 25%; **Figure 1H**), indicating that threat exposure information is encoded by the activity pattern of at least a subset of AHA^VGAT^ neurons. Remarkably, we observed an increase in AHA^PV^ neuronal activity during threat exposure (*n* = 13 neurons from 2 mice), which multiple comparisons testing revealed was significant compared to post-threat stages (**Figure 1I–K**). Predictions of mouse behavioral stage from AHA^PV^ neuronal activity using a quadratic SVM were well above chance accuracy (75.0% compared to 25%; **Figure 1L**). With the exception of cubic KNN for AHA^PV^ neurons, all computational models tested were capable of predicting behavioral stage from the activity patterns of both AHA^VGAT^ and AHA^PV^ neurons (**Supplementary Table 1**).

**Figure 1.**
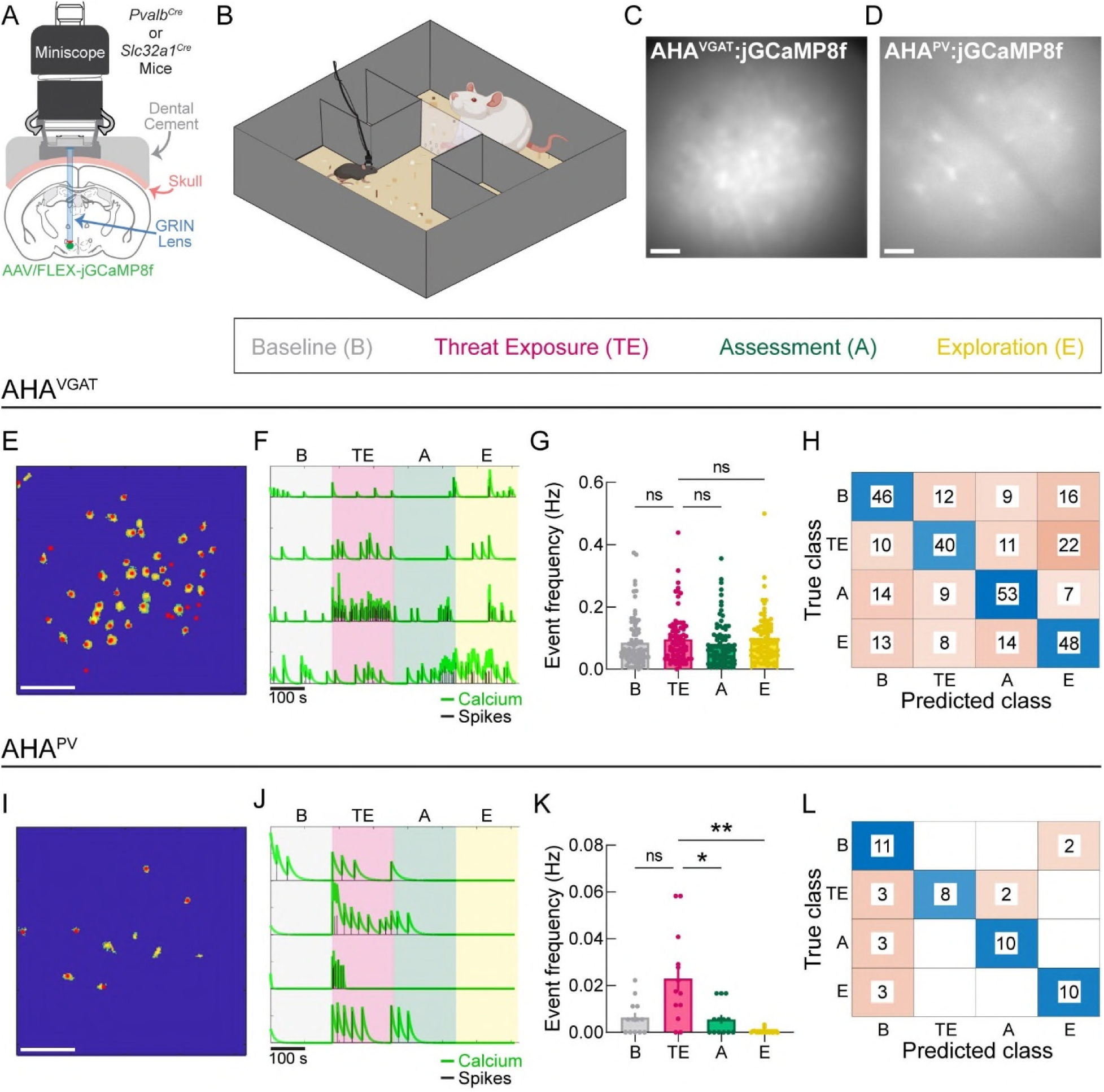
Exposure to a predatory threat triggers an increase in AHA^PV^ neuronal activity. **(A)** Representative diagram of miniscope with GRIN lens implantation dorsal to the AHA. **(B)** Schematic of the predatory threat assay showing the H-box arena with a rat in the top compartment and an experimental mouse in the bottom compartment separated by a transparent wall-blocked hallway. **(C−D)** Representative maximum intensity projections of *in vivo* functional images of **(C)** AHA^VGAT^ neurons (*n* = 83 from 3 mice) and **(D)** AHA^PV^ neurons (*n* = 13 cells from 2 mice). Scale bars = 500 µm. **(E)** Spatial footprints of AHA^VGAT^ neurons. **(F)** Representative deconvolution for extracted calcium signal (green) and estimated periods of neuronal firing (“spikes”; black) from AHA^VGAT^ neurons during baseline (B; gray region), mouse exposure to a rat (TE; magenta region), apparatus assessment after rat removal (A; green region), and exploration of the rat zone after wall removal (E; yellow region). **(G)** AHA^VGAT^ neuron event frequency during each stage of the predatory threat assay. **(H)** Confusion matrix showing quadratic SVM classification performance when predicting behavioral stage based on AHA^VGAT^ neuron activity during the predatory threat test (Accuracy = 56.8%). **(I)** Spatial footprints of AHA^PV^ neurons. **(J)** Representative deconvolution for extracted calcium signal (green) and spike detection (black) from AHA^PV^ neurons during baseline (B; gray region), mouse threat exposure to a rat (TE; magenta region), mouse apparatus assessment after rat removal (A; green region), and mouse exploration of the rat zone after wall removal (E; yellow region). **(K)** AHA^PV^ neuron event frequency during each stage of the predatory threat assay. **(L)** Confusion matrix showing quadratic SVM classification performance when predicting behavioral stage based on AHA^PV^ neuron activity during the predatory threat test (Accuracy = 75.0%). Data are represented as mean ± S.E.M. \**p* < 0.05; \*\**p* < 0.01; ns, not significant.

### AHA^PV^ neurons are fast-spiking and express ion channels implicated in setting and regulating fast-spiking activity

It has been widely demonstrated that PV-positive neurons across the brain exhibit a fast-spiking phenotype (Bartholome et al., 2020; Markram et al., 2004; Siemian et al., 2020). Therefore, we characterized the electrophysiological properties of AHA^PV^ neurons and determined the expression of specific ion channels in these neurons by using brain slice electrophysiology and fluorescent *in situ* hybridization, respectively, to determine whether AHA^PV^ neurons exhibit the characteristic hallmarks of other PV neuron populations.

We performed whole-cell patch clamp recordings in brain slices from *Pvalb^Cre/+^;Rosa26^LSL-tdtomato/LSL-tdTomato^* mice and found that these neurons fire action potentials at high-frequency (111.07 ± 7.51 Hz; *n* = 30 neurons; **Figure 2A and Table 1**). Moreover, AHA^PV^ electrophysiological characteristics (**Table 1**) are similar to the properties of other PV-positive neurons throughout the brain (DeFelipe et al., 2013; Hu et al., 2014; Kisner *et al*., 2018).

**Figure 2.**
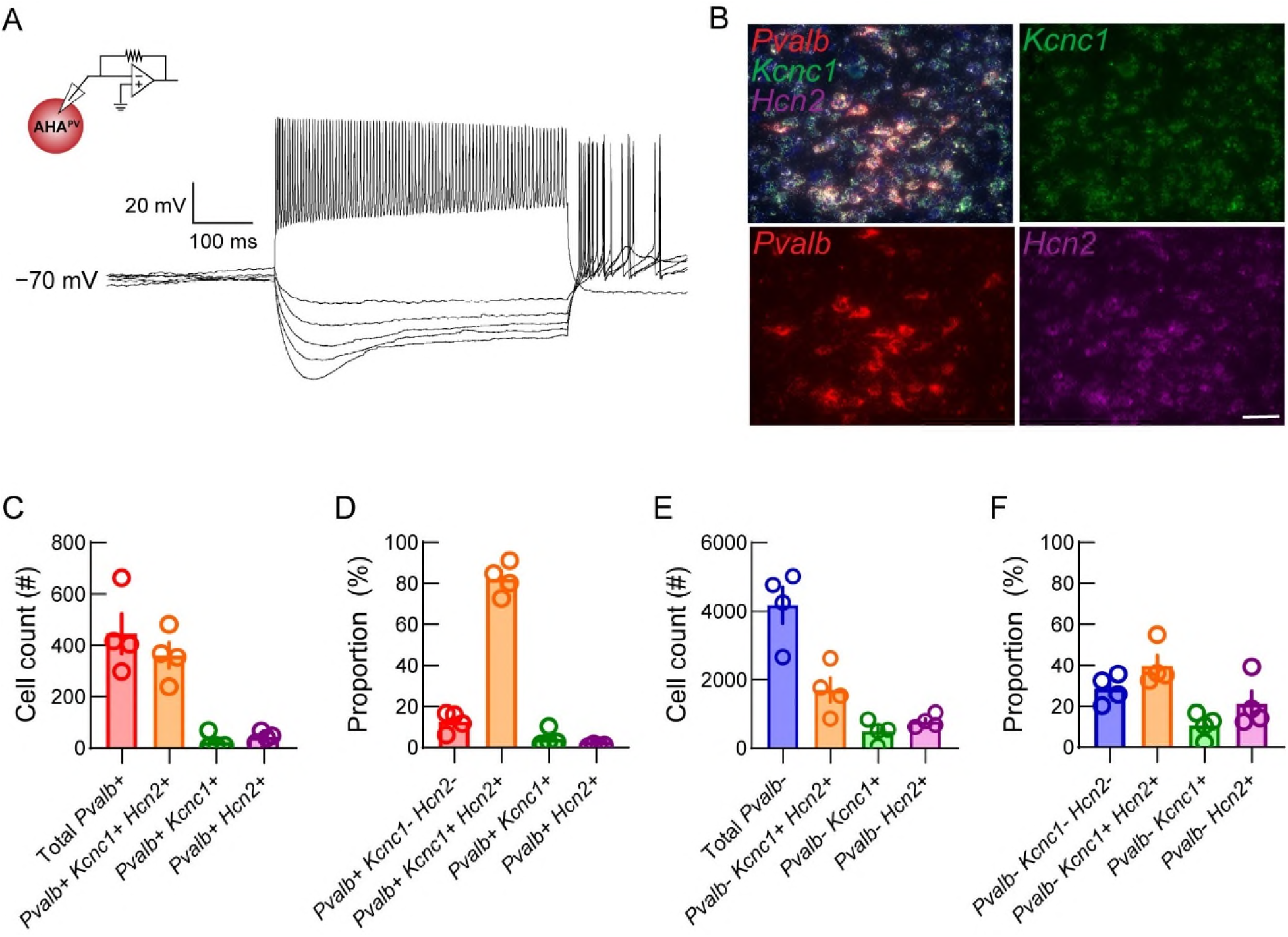
AHA^PV^ neurons are fast-spiking and express ion channels correlated with a fast-spiking phenotype. **(A)** Representative current clamp recording of hyperpolarization and fast-spiking in an AHA^PV^ neuron from a *Pvalb^Cre^;Rosa26^LSLtdTomato^* transgenic mouse. **(B)** Representative images of a coronal section (40x) depicting composite with DAPI (*top left*) and individual channel images of fluorescent *in situ* hybridization for *Kcnc1* (green), *Pvalb* (red), and *Hcn2* (magenta) mRNA. Scale bar = 200 µm. **(C−D)** Sum of cell counts (*n* = 4 mice) in **(C)** raw and **(D)** proportion format for cellular co-localization of *Pvalb^+^* neurons with *Kcnc1* and *Hcn2* mRNA. **(E−F)** Cell counts in **(E)** raw and **(F)** proportion format for cellular co-localization for parvalbumin mRNA-negative neurons with *Kcnc1* and *Hcn2* mRNA. Data are represented as mean ± S.E.M.

**Table 1.**
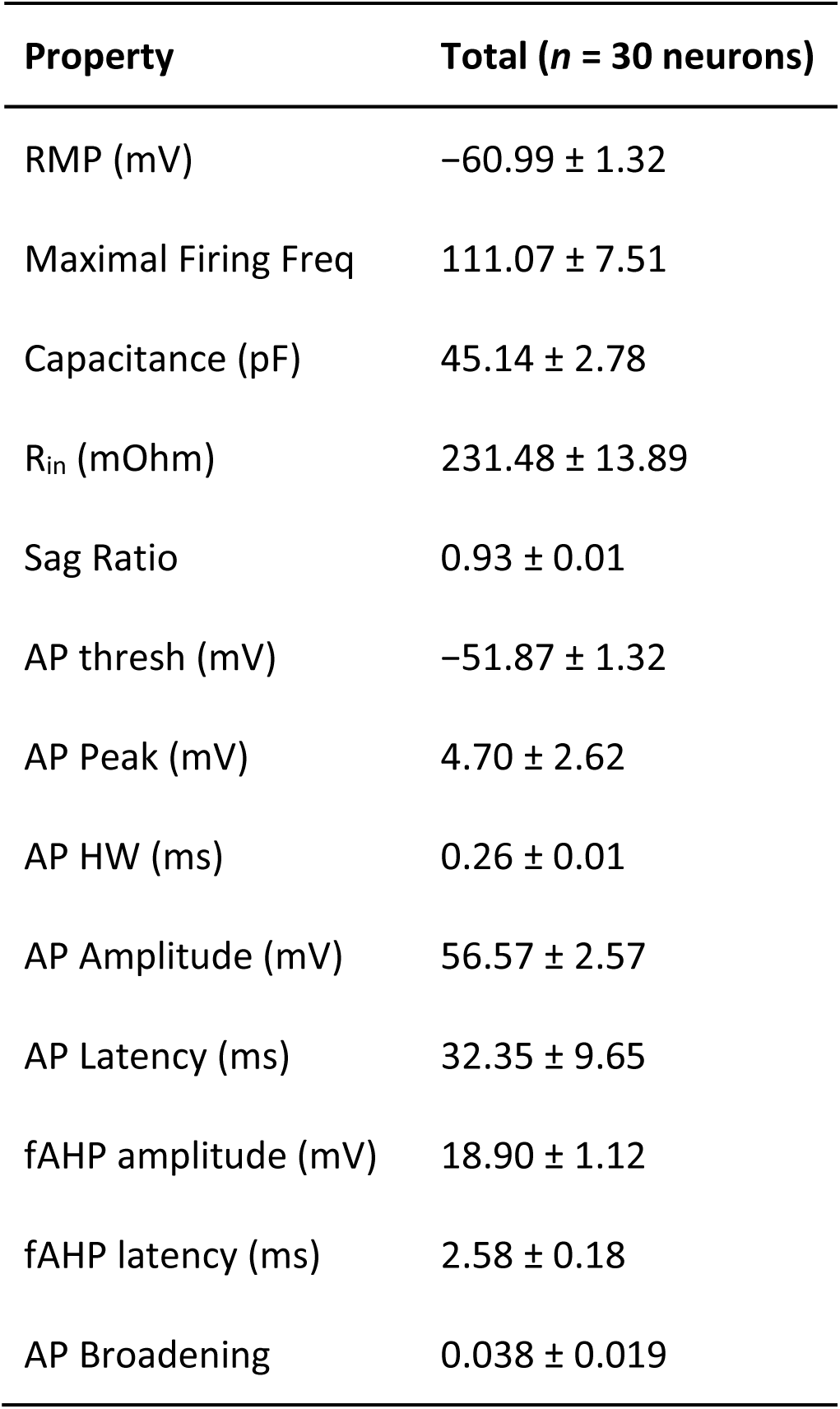
Electrophysiological properties of AHA^PV^ neurons.

| Property | Total (*n* = 30 neurons) |
| --- | --- |
| RMP (mV) | -60.99 ± 1.32 |
| Maximal Firing Freq | 111.07 ± 7.51 |
| Capacitance (pF) | 45.14 ± 2.78 |
| R<sub>in</sub> (mOhm) | 231.48 ± 13.89 |
| Sag Ratio | 0.93 ± 0.01 |
| AP thresh (mV) | -51.87 ± 1.32 |
| AP Peak (mV) | 4.70 ± 2.62 |
| AP HW (ms) | 0.26 ± 0.01 |
| AP Amplitude (mV) | 56.57 ± 2.57 |
| AP Latency (ms) | 32.35 ± 9.65 |
| fAHP amplitude (mV) | 18.90 ± 1.12 |
| fAHP latency (ms) | 2.58 ± 0.18 |
| AP Broadening | 0.038 ± 0.019 |

Fast-spiking characteristics are correlated with the expression of specific ion channels such as the delayed rectifier voltage-gated potassium channel KCNC1 (Kv3.1) (Du et al., 1996; Rosato-Siri et al., 2015) and the potassium/sodium hyperpolarization-activated cyclic nucleotide-gated ion channel 2 (HCN2) (Aponte et al., 2006). Thus, to determine whether these molecular markers are expressed in AHA^PV^ neurons, we performed fluorescent *in situ* hybridization on brain slices containing the AHA (*n* = 4 mice; **Figure 2B**) and analyzed these data using Autocount, a custom ImageJ macro tool (Laing, 2022). We observed that AHA^PV^ neurons express both *Kcnc1* and *Hcn2* mRNA (360.3 ± 49.49 neurons of 445 ± 77.04 total *Pvalb^+^* neurons; **Figure 2C**). Moreover, we found that proportionally the vast majority of neurons expressed all three transcripts (*Pvalb^+^ Kcnc1^+^ Hcn2^+^*, 82.20% ± 3.885; **Figure 2D**). Furthermore, we observed that few *Pvalb* mRNA*-*negative cells expressed *Kcnc1* and *Hcn2* (1696 ± 361.5 neurons of 4171 ± 529.0 total *Pvalb^−^* neurons; **Figure 2E**), which represents about one third of this neuronal population (33.17% ± 3.885; **Figure 2F**), indicating heterogeneity among cells that do not express *Pvalb* mRNA in the AHA. Chi-square analysis of the proportion of AHA^PV^ neurons versus *Pvalb* mRNA*-* negative cells revealed significantly higher colocalization of *Hcn2* and *Kcnc1* mRNA in AHA^PV^ neurons (χ^2^ = 93.79, df = 3). Together, our results demonstrate that AHA^PV^ neurons express key ion channels for a fast-spiking phenotype and further support the idea that the AHA is composed of a diverse collection of cell types.

### AHA^PV^ neurons form functional excitatory synapses onto PMD neurons

Next, we mapped the axonal projections of AHA^PV^ neurons by unilaterally injecting the AHA of *Pvalb^Cre^* mice with a Cre-dependent AAV expressing the presynaptic protein synaptophysin fused to the fluorophore mCherry (Opland et al., 2013). Using this strategy, we identified the premammillary dorsal nucleus (PMD) as a postsynaptic target of AHA^PV^ neurons (**Figure 3A–B**, *n* = 10 mice). Additionally, we unilaterally injected the retrograde tracer Fluoro-Gold into the PMD of *Pvalb^Cre^;Rosa26^LSLtdTomato^*transgenic mice (**Figure 3C–D**, *n* = 4 mice). Analysis of Fluoro-Gold colocalization with tdTomato-positive AHA^PV^ neurons showed that more than half of the total AHA^PV^ population (66.4% ± 5.4) projects to the PMD. Together, these results suggest that AHA^PV^ axonal projections to the PMD (AHA^PV^→PMD) could potentially regulate neuronal activity within the PMD to trigger behavior.

**Figure 3.**
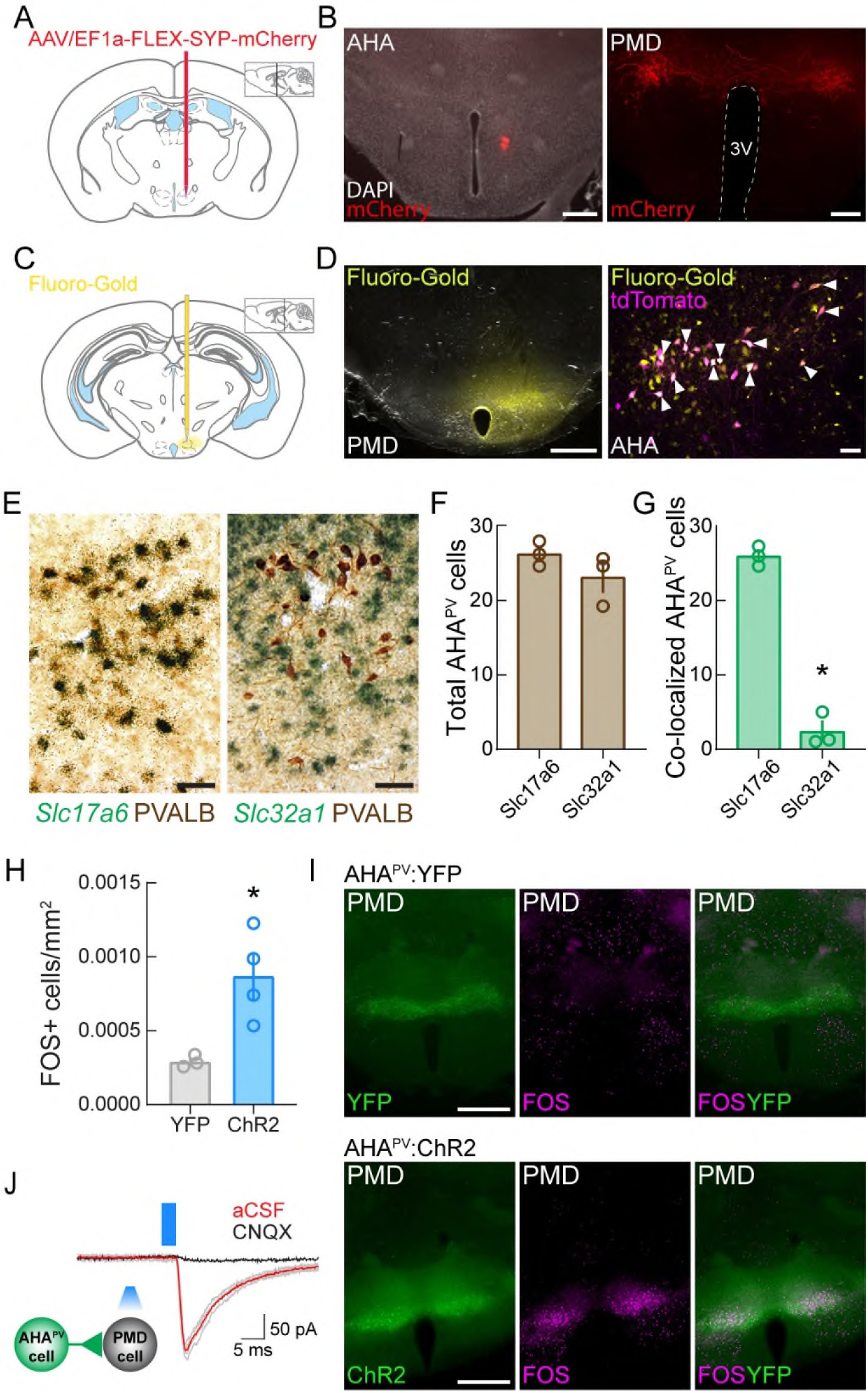
AHA^PV^ neurons are glutamatergic and provide excitatory inputs onto PMD neuron. **(A)** Schematic depicting unilateral injection of the anterograde tracer AAV/EF1a-FLEX-SYP-mCherry into the AHA of *Pvalb^Cre^* mice. **(B)** Representative images showing viral transduction in AHA^PV^ neurons (*left panel*; Scale bar = 500 µm) and axonal projections in the dorsal premammillary nucleus (PMD; *right panel*; Scale bar = 200 µm). **(C)** Schematic depicting unilateral Fluoro-Gold injections into the PMD of *Pvalb^Cre^;Rosa26^LSLtdTomato^*transgenic mice to determine specific inputs from AHA^PV^ neurons. **(D)** Representative images showing Fluoro-Gold at the PMD injection site (*left panel*; Scale bar = 500 µm) and co-localization of Fluoro-Gold and tdTomato in AHA^PV^ neurons (*right panel*; Scale bar = 50 µm). **(E)** Representative photomicrographs of coronal sections showing that parvalbumin (PVALB) immunoreactive cells (brown) in the AHA primarily co-localize with *Vglut2* mRNA but not *Vgat* mRNA (green grain aggregates), Scale bars = 50 µm. **(F)** There were no significant differences in the total number of AHA^PV^ neurons counted between matched coronal slices**. (G)** AHA^PV^ neurons predominantly co-localize with the glutamatergic marker compared to the GABAergic marker. **(H)** Number of FOS-expressing cells per unit area in the PMD in AHA^PV^:YFP (*n* = 3 mice) and AHA^PV^:ChR2 groups (*n* = 4 mice) after photoactivation of AHA^PV^ neurons. **(I)** Representative images showing AHA^PV^:YFP axons (green) and FOS (magenta) immunostaining in the PMD (*top panels*; Scale bar = 500 µm) and AHA^PV^:ChR2 axons (green) and FOS (magenta) immunostaining in the PMD (*bottom panels*; Scale bar = 500 µm). **(J)** Representative traces of excitatory postsynaptic currents in gray (EPSCs; *n* = 8/18 neurons) evoked on neurons within the PMD by photostimulation of AHA^PV^:ChR2 axonal projections (3- ms light pulses; 470 nm LED), these EPSCs were blocked by bath application of the AMPA receptor antagonist CNQX (black trace; *n* = 3 neurons). Red and black traces are the average of 5 consecutive sweeps. Data are represented as mean ± S.E.M. \**p* < 0.05.

To determine the profile of neurotransmitters expressed by AHA^PV^ neurons, we performed *in situ* hybridization for molecular markers associated with glutamatergic (*Slc17a6*) and GABAergic transmission (*Slc32a1*) (**Figure 3E**). While there were no differences in the total number of AHA^PV^ neurons counted per group (**Figure 3F**, *n* = 3 mice per group, unpaired two-tailed Welch’s t-test, *t*_1.48_ = 0.2434), there were significantly more AHA^PV^ neurons colocalized with the glutamatergic marker (26.0 ± 1.33 neurons) than the GABAergic marker (2.41 ± 2.24 neurons), indicating that these neurons are predominantly glutamatergic (**Figure 3G**, unpaired two-tailed Welch’s *t*-test, *t*_3.25_ = 0.0004).

We next sought to investigate changes in PMD neuronal activity triggered by photoactivation of AHA^PV^ neurons. For this, we photoactivated (20 Hz, 10−15 mW, 10 ms) AHA^PV^:ChR2 and AHA^PV^:YFP mice for one hour prior to euthanasia (*n* = 4 and *n* = 3, respectively). Using expression of FOS as an indirect marker for neuronal activity (Hudson, 2018), we counted the number of FOS-positive cells in the PMD and compared them between the AHA^PV^:ChR2 and AHA^PV^:YFP groups. We found that FOS expression is significantly elevated in the PMD after AHA^PV^ neuronal activation (**Figure 3H–I**, Welch’s two-tailed *t*-test, *t*_3.15_ = 3.814, *p* = 0.0291). We then performed channelrhodopsin-2-assisted circuit mapping (CRACM) to determine whether AHA^PV^ neurons are synaptically connected to PMD neurons. For this, we bilaterally injected a Cre-dependent AAV expressing ChR2-YFP into the AHA of *Pvalb^Cre^*mice and conducted whole-cell recordings of individual cells in the PMD. CRACM revealed that AHA^PV^ neurons release the excitatory neurotransmitter glutamate and provide excitatory inputs onto PMD neurons (**Figure 3J,** *n* = 8/18 neurons, mean amplitude = −90.34 ± 28.10 pA, mean latency = 5.84 ± 0.25 ms). Additionally, the AMPA receptor antagonist CNQX was bath applied to block light-evoked excitatory postsynaptic currents (EPSCs) in PMD cells (*n* = 3; EPSC_aCSF_ mean = −100.9 pA ± 41.73, EPSC_CNQX_ mean = −7.56 pA ± 1.33), and we observed a significantly reduced amplitude in CNQX when normalized to the mean amplitude recorded during optically evoked currents in aCSF (*n* = 3, Welch’s paired two-tailed *t*-test, *t_2_* = 23.07, *p* = 0.019). Together, these results demonstrate that AHA^PV^ neurons provide glutamatergic input to the PMD to promote activation of neuronal circuits within the PMD.

### Photoactivation of AHA^PV^ neurons is aversive and triggers escape responding

We conducted several behavioral tests to determine the effects of optogenetic manipulations of AHA^PV^ neurons. To specifically target these neurons, we bilaterally injected the AHA of *Pvalb^Cre^* mice with Cre recombinase-dependent viral vectors for inhibition (iC++:YFP; *n* = 9 AHA^PV^:iC++ mice), activation (ChR2:YFP; *n = 8* AHA^PV^:ChR2 mice for somatic stimulation; *n = 7* AHA^PV^:ChR2→PMD mice for axonal stimulation), or fluorophore control (YFP; *n* = 11 AHA^PV^:YFP mice). A four-group design was used to assess behavioral responses during inhibition and somatic/axonal activation compared to controls. Three groups were bilaterally implanted with optical fibers dorsal to the AHA, and one group was bilaterally implanted with optical fibers dorsal to the PMD to photostimulate AHA^PV^ axonal projections in this region (**Figure 4A**). Post-mortem histological analysis was conducted to map viral transduction (**Figure 4B**).

**Figure 4.**
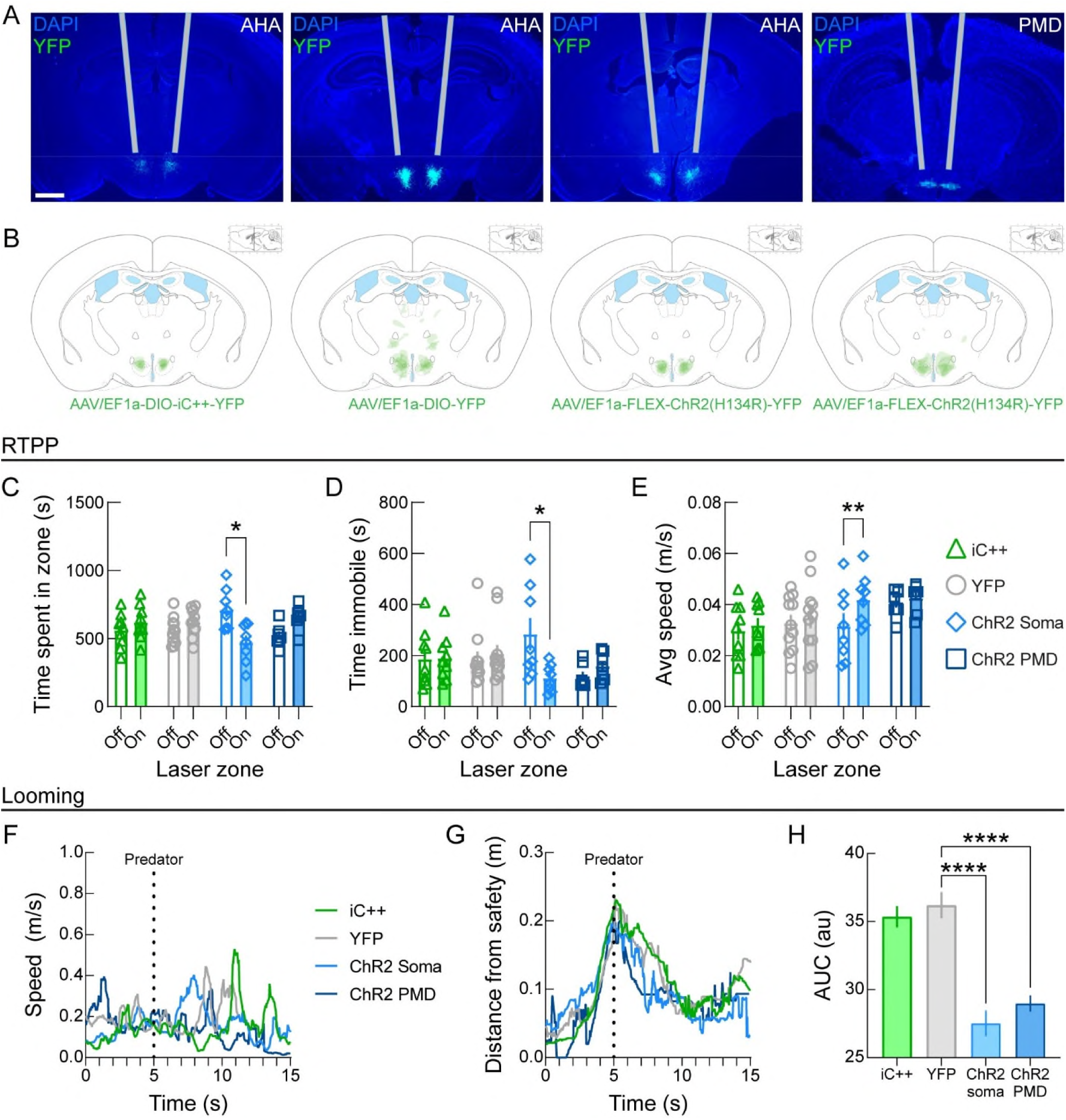
Photoactivation of AHA^PV^ neurons or the AHA^PV^→PMD pathway promotes escape behavior. **(A)** Representative images of AHA^PV^ soma or terminal transduction for the AHA^PV^:iC++, AHA^PV^:YFP control, AHA^PV^:ChR2 soma, and AHA^PV^:ChR2→PMD groups with estimated optical fiber paths (Scale bar = 1 mm). **(B)** Diagrams depicting viral transduction in *Pvalb^Cre^* mice for AHA^PV^:iC++, AHA^PV^:YFP control, AHA^PV^:ChR2 soma, and AHA^PV^:ChR2→PMD groups. Transduced areas were traced from overlaid coronal sections (20% opacity per mouse) onto each diagram. **(C−E)** Optogenetic manipulation of AHA^PV^ neurons during real-time place preference reveals **(C)** significant avoidance of the Laser-On zone by the AHA^PV^:ChR2 soma group as well as **(D)** decreased time spent immobile and **(E)** increased speed within the laser-paired side of the apparatus. (AHA^PV^:iC++ *n* = 9, AHA^PV^:YFP control *n* = 11, AHA^PV^:ChR2 soma *n* = 8, and AHA^PV^:ChR2→PMD *n* = 7). **(F−H)** Optogenetic manipulation of AHA^PV^ neurons during the simulated looming predator assay reveals an alteration of flight behavior reflected in **(F)** a left-shifted speed profile for AHA^PV^:ChR2 soma mice and **(G)** a rapid reduction in distance from safety in the AHA^PV^:ChR2 soma and AHA^PV^:ChR2→PMD groups, which is seen as **(H)** a reduced area under the curve (AUC) in both of these groups. (AHA^PV^:iC++ *n* = 9, AHA^PV^:YFP *n* = 11, AHA^PV^:ChR2 *n* = 8, AHA^PV^:ChR2→PMD *n* = 5). Data are represented as mean ± S.E.M. \**p* < 0.05; \*\**p* < 0.01; \*\*\*\**p* < 0.0001.

We first used a real-time place preference (RTPP) assay to measure the effects of AHA^PV^ neuron manipulation on avoidance. During this assay, one side of the testing arena was paired with photoactivation or photoinhibition depending on the cohort, and mice were permitted to freely move throughout both sides of the arena (i.e., Laser-Off and Laser-On). We observed a significant time × transgene effect (**Figure 4C**, Two-way RM ANOVA, *F*_3, 31_ = 4.174, *p* = 0.0136) and multiple comparisons revealed a significant reduction in time spent in the Laser-On zone specifically for the AHA^PV^:ChR2 mice (*p* = 0.0247). Further analysis showed that there was a significant stimulation × transgene effect on time immobile (**Figure 4D,** Two-way RM ANOVA, *F*_3, 31_ = 3.727, *p* = 0.0383). Notably, the AHA^PV^:ChR2 group was the only group to exhibit a significant increase in time immobile in the Laser-Off zone compared to the Laser-On zone (*p* = 0.0128). Significant stimulation effects on average speed in the Laser-On zone were also detected (**Figure 4E**, Two-way RM ANOVA, *F*_1, 31_ = 8.340, *p* = 0.0070) with stimulation × transgene effects just above significance level (Two-way RM ANOVA, *F*_3, 31_ = 2.153, *p* = 0.1136). Multiple comparisons revealed a significant within group effect exclusively in the AHA^PV^:ChR2 somatic stimulation group (*p* = 0.0046). Together, these results demonstrate that somatic activation of AHA^PV^ neurons is aversive and triggers locomotive avoidance.

Because we observed a significant increase in AHA^PV^ neuronal activity during exposure to a rat, we next examined the effects AHA^PV^ neuron manipulations during the ethologically relevant predator-looming task (Yilmaz and Meister, 2013). During this task, a mouse is placed in an arena with a covered nest representing a “safe zone.” After the mouse claims the nest, a round black circle (looming stimulus) travels across an overhead monitor to simulate a predator. Photoactivation or photoinhibition occurred at the onset of looming stimulus delivery. We found that speed profiles were not affected by optogenetic manipulations of AHA^PV^ neurons or the AHA^PV^→PMD pathway **(Figure 4F)**. However, distance from the safety zone estimated as the area under the curve (AUC) was significantly altered between groups (**Figure 4G**, One-way ANOVA, *F*_3, 23_ = 25.21, *p* < 0.0001). Multiple comparisons testing revealed that both AHA^PV^:ChR2 and AHA^PV^:ChR2→PMD groups displayed a significant reduction in distance from the safety zone compared to the AHA^PV^:YFP and AHA^PV^:iC++ groups (*p* < 0.0001), but not between each other **(Figure 4H)**. Thus, these results demonstrate that activation of both AHA^PV^ neurons and the AHA^PV^→PMD pathway evokes escape responding.

### Cell-type specific ablation of AHA^PV^ neurons does not affect anxiety-like behavior but impairs escape responding

We sought to determine the effects of AHA^PV^ neuron ablation on behavior by bilaterally injecting a Cre-dependent viral vector driving the expression of caspase-3 (CASP3) (Yang et al., 2013) or the control fluorophore mCherry into the AHA of *Pvalb^Cre^* mice (**Figure 5A**). Representative images of parvalbumin immunostaining for each group depict the presence of parvalbumin-positive neurons in control AHA^PV^:mCherry mice and genetic ablation of these neurons in AHA^PV^:Casp3 mice (**Figure 5B**). Quantitative analysis revealed a significant reduction in the number of parvalbumin-positives neurons in the AHA^PV^:Casp3 mice (**Figure 5C**, AHA^PV^:mCherry control group, mean: 62.00 ± 8.55 neurons; AHA^PV^:Casp3 group, mean: 18.75 ± 2.99 neurons; Welch’s two-tailed *t*-test *t*_3.892_ = 3.892, *p* = 0.0017). Subsequently, we used the open field test to assess behavior in a novel environment and did not observe significant differences between the AHA^PV^:Casp3 mice compared to the AHA^PV^:mCherry controls in time spent in the center zone of the arena (**Figure 5D**, Two-tailed *t*-test, *t*_13_ = 0.6404, *p* = 0.5331) or mean speed in center zone (**Figure 5E**, Two-tailed *t*-test, *t*_13_ = 0.1184, *p* = 0.9076). To further examine the effects of AHA^PV^ neuron ablation on anxiety-like behaviors, we used both a light-dark transition test and the elevated plus maze. No significant differences between groups were observed on time spent in the light zone (**Figure 5F**, Two-tailed *t*-test, *t*_13_ = 0.3363, *p* = 0.7420) or time spent in the open arms of the elevated plus maze (**Figure 5G**, Two-tailed *t*-test, *t*_13_ = 0.5565, *p* = 0.5873), suggesting that in this context, AHA^PV^ neuronal activity does not seem necessary for regulating anxiety-like behavior.

**Figure 5.**
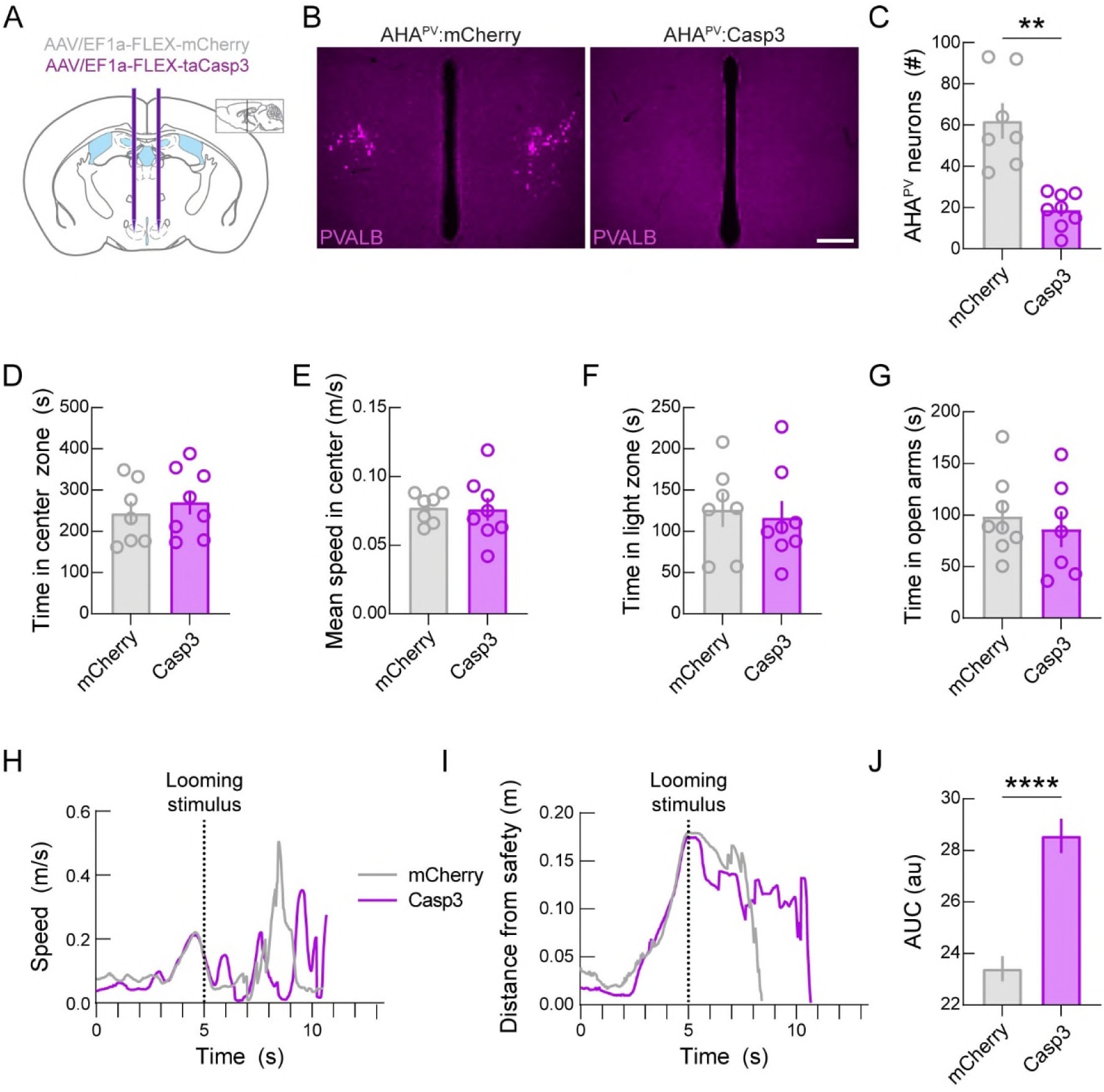
Genetic ablation of AHA^PV^ neurons impairs escape responding during a predator- looming task. **(A)** Diagram depicting bilateral injection of mCherry control and caspase (Casp3) viruses into the AHA of *Pvalb^Cre^* mice. **(B)** Representative parvalbumin immunostaining (PVALB, magenta) in the AHA of AHA^PV^:mCherry control (*n* = 7) and AHA^PV^:Casp3 mice (*n* = 8). **(C)** Quantification shows a significant reduction of PVALB neurons in AHA^PV^:Casp3 mice compared to AHA^PV^:mCherry controls. **(D−E)** No significant effects were observed during the open field test for **(D)** time in center zone or **(E)** mean speed in center zone. **(F)** No significant effects were detected for time in the light zone during the light-dark transition test. **(G)** No changes were detected for time spent in the open arms of the elevated plus maze. **(H)** In the predator-looming test, escape response is impaired in AHA^PV^:Casp3 mice compared to AHA^PV^:mCherry controls. **(I)** Arrival to the safety nest is delayed in AHA^PV^:Casp3 mice compared to AHA^PV^:mCherry controls. **(J)** Quantification of area under the curve (AUC) by group indicates a significantly larger AUC for the AHA^PV^:Casp3 group compared to controls. Data are represented as mean ± S.E.M. \*\**p* < 0.01; \*\*\*\**p* < 0.0001.

Next, we investigated the effects of AHA^PV^ neuron ablation on escape behavior using the predator-looming task. We did not observe significant differences between groups in speed profiles (**Figure 5H**). Notably, the distance from the safety zone was right-shifted for the AHA^PV^:taCasp3 group compared to controls (**Figure 5I**). Further AUC analysis revealed that AHA^PV^ neuron ablation evoked a significant increase in the distance from the safety zone in the AHA^PV^:Casp3 mice compared to controls (**Figure 5J**, Two-tailed *t*-test, *t*_12_ = 0.1485, *p* = 0.8844) indicating that ablation of these neurons significantly impairs escape responding.

### Changes in neuronal activity of downstream brain regions triggered by photoactivation of AHA^PV^ neurons

Finally, we examined brain-wide discrete changes in neuronal activity during AHA^PV^ photoactivation. For this, we compared [^18^F]fluorodeoxyglucose (FDG)-PET scans (Boehm et al., 2021) between AHA^PV^:ChR2 and AHA^PV^:tdTomato mice (*n* = 4 per group). Because the brain uses glucose as its main energy substrate, FDG is taken up by active neurons (Pacák et al., 1969; Sokoloff et al., 1977), and brain activity patterns can be determined by monitoring the accumulation of FDG in different brain regions. By pairing FDG-PET with cell type-specific neuromodulation approaches such as optogenetics (Thanos et al., 2013a), regional changes in neuronal activity across the entire brain triggered by modulation of a specific neuron population can be determined (Michaelides and Hurd, 2015).

We unilaterally injected *Pvalb^Cre^* mice with Cre-dependent ChR2-tdTomato or tdTomato (control) viral vectors into the AHA and implanted optical fibers unilaterally above this area (**Figure 6A–B**). Anesthetized mice were placed into a PET/computed tomography (CT) scanner and injected with FDG followed by alternating periods of photostimulation-off and -on (**Figure 6C**). We found that photoactivation of AHA^PV^ neurons evoked significant changes in FDG uptake across multiple brain regions as indicated by *t*-values of between groups comparisons (**Figure 6D**). Specifically, significant increases in brain activity were observed in the olfactory tubercle ipsilateral to the photostimulated hemisphere (bregma +1.5), while significant decreases were detected in the contralateral ventral striatum and amygdala (bregma +0.5 to −1.0). In addition, significantly increased activity was observed in the ipsilateral retrosplenial cortex and amygdala (bregma −2.0), ipsilateral substantia nigra reticulata (bregma −3.0 to bregma −4.0), contralateral subiculum (bregma −3.0 to bregma −4.0), and ipsilateral cochlear nucleus (bregma −5.0). Together, these results indicate that activation of AHA^PV^ neurons evokes discrete activation of downstream areas involved in arousal, affective, and defensive behaviors such as the amygdala and the substantia nigra.

**Figure 6.**
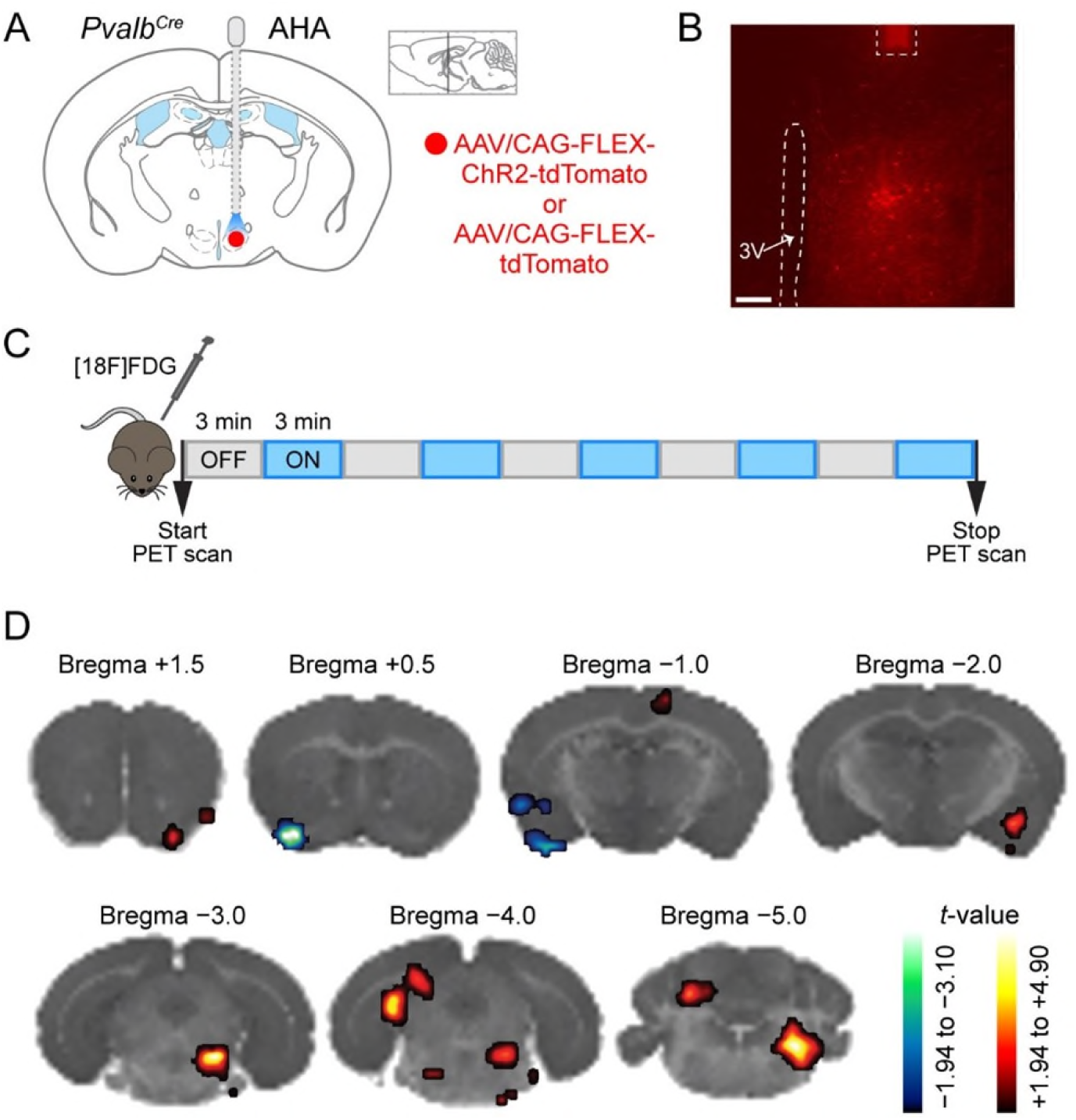
PET scans during photoactivation of AHA^PV^ neurons reveal changes in activity in brain regions involved in defensive behaviors. **(A)** Schematic of unilateral viral injection into the AHA of *Pvalb^Cre^* mice with an optical fiber implanted dorsal to the injection site. **(B)** Representative image showing fiber optic placement (dotted line) over a cluster of AHA^PV^ neurons. 3V, third ventricle. Scale bar = 200 µm. **(C)** Diagram depicting PET scan experiment. Scanning started immediately after FDG injection (i.p.) into anesthetized mice and spanned five OFF/ON photostimulation epochs (10-ms pulse, 20 Hz). **(D)** Changes in brain activity were detected in several brain regions such as increases in the olfactory tubercle ipsilateral to the photostimulated hemisphere (bregma +1.5), decreases in the contralateral ventral striatum and amygdala (bregma +0.5 to −1.0); also increases in the ipsilateral retrosplenial cortex and amygdala (bregma −2.0), ipsilateral substantia nigra reticulata (bregma −3.0 to bregma −4.0), contralateral subiculum (bregma −3.0 to bregma −4.0), and ipsilateral cochlear (bregma −5.0). Color bars represent increasing and decreasing *t* values.

## DISCUSSION

Previous studies have elegantly demonstrated that activation of excitatory inputs to the AHA triggers defensive-like behavior such as immobility (Wang *et al*., 2015). In addition, escaping and freezing have been reported upon stimulation of glutamatergic receptors in the AHA (Falconi-Sobrinho and Coimbra, 2018). Moreover, administration of the cholinergic agonist carbachol in the AHA increases the duration of immobile episodes in a muscarinic receptor dependent fashion (de Oliveira et al., 1997). While these approaches yield significant comparative insight into the function of the AHA, the specific roles of molecularly defined neuron populations within this region have yet to be determined. Here, we characterized a cluster of AHA^PV^ neurons that are fast-spiking, glutamatergic, and send axonal projections to the PMD, a brain region involved in flexible escape responses (Wang et al., 2021).

Our electrophysiological analyses showed that some of the properties of AHA^PV^ neurons are consistent with those of other glutamatergic parvalbumin neurons throughout the brain (Siemian *et al*., 2020). *In situ* hybridization and electrophysiological recordings revealed that these neurons are predominantly glutamatergic and their fast-spiking characteristic is comparable to other glutamatergic parvalbumin neurons elsewhere in the brain such as the entopeduncular nucleus (Wallace et al., 2017) and the lateral hypothalamus (Kisner *et al*., 2018; Siemian *et al*., 2019). This is distinct from glutamatergic parvalbumin-expressing neurons in the ventral pallidum, which were reported to fire at a mean maximum of 25 Hz (Tooley et al., 2018).

In terms of behavior, our findings are consistent with previous studies showing that VMH→AHA stimulation evokes defense-like motor responses characterized by decreased immobility and increased locomotor activity (Wang *et al*., 2015). First, we showed that AHA^PV^ neuronal activity significantly increases when a mouse is exposed to a rat—a predatory threat— demonstrating that these neurons are modulated by threat-related stimuli. Moreover, using machine learning models, we were able to predict the behavioral stage of the mouse based on the activity patterns of AHA^VGAT^ and AHA^PV^ triggered by the predatory threat.

During our real-time place preference experiments, we found that photoactivation of AHA^PV^ neurons is aversive and triggers locomotive avoidance. However, we did not observe significant changes on such responses when photostimulating the AHA^PV^→PMD pathway. Thus, it is possible that this pathway encodes more complex aspects of escape behaviors rather than stereotyped fear-related escape. Our results on the predator-looming task support this idea, as a mouse’s return to a safety zone is promoted by photoactivation of both AHA^PV^ neurons and the AHA^PV^→PMD pathway. Thus, escape responding seems to be modulated by the activity of AHA^PV^ neurons and the AHA^PV^→PMD pathway. Notably, photoinhibition of AHA^PV^ neurons did not affect escape responding during the predator-looming assay. However, escape responding was impaired after specific ablation of AHA^PV^ neurons, indicating that long-term loss of AHA^PV^ neuronal activity attenuates returning to a safety zone. This is consistent with previous studies showing decreased autonomic/cardiovascular flexibility in mice lacking AHA^PV^ neurons (Mittag *et al*., 2013). While the rationale for the discrepancy observed during the predator-looming assay between acute and chronic loss of AHA^PV^ neuron function has yet to be determined, this could be attributed to long-term adaptations of sympathetic drive associated with the absence of AHA^PV^ neurons. Interestingly, anxiety-like behaviors measured by the open field and elevated plus maze assays do not seem to be affected by AHA^PV^ neuron ablation. Together, these results further support that these neurons may orchestrate more complex aspects of escape behaviors.

Our anterograde and retrograde tracing experiments revealed the PMD as a post-synaptic target of AHA^PV^ neurons. These results were further supported by our CRACM experiments demonstrating that AHA^PV^ neurons are synaptically connected to neurons within the PMD. Our data showed that photoactivation of the AHA^PV^→PMD pathway promotes escape responding during exposure to a predatory stimulus. Previous studies have demonstrated that the PMD is activated in response to innate fear (Mendes-Gomes et al., 2020) as well as social confrontation (Motta et al., 2009). Moreover, the main inputs to the PMD are from the AHA (Canteras et al., 2001; Comoli et al., 2000). Furthermore, the PMD becomes active in rats exposed to threat or stressful conditions (Kim et al., 2017; Mendes-Gomes *et al*., 2020), and lesion of the PMD abolishes locomotor arousal defense in response to predator odors (Canteras et al., 1997). More complex and ethologically relevant behavioral assays will reveal whether AHA^PV^ neurons encode passive components of defensive behavior, such as hiding and scanning a risky environment.

Whole-brain metabolic mapping using FDG-PET revealed several downstream brain regions modulated by photoactivation of AHA^PV^ neurons. Regions related to sensory processing showed increased activity, including the olfactory tubercle and cochlear nucleus. The retrosplenial cortex showed significantly elevated activity on the ipsilateral side of the photostimulated hemisphere, suggesting potential effects on perceptual and memory functions related to contextual cues (Kwapis et al., 2015) that induce fear (Corcoran et al., 2011). We also observed significantly increased activity in the substantia nigra reticulata (SNr) which is consistent with previous studies showing the involvement of the SNr in escape-mediated behavior (Coimbra et al., 1989). Moreover, the ventral striatum and the amygdala are also brain regions activated by photoactivation of AHA^PV^ neurons. Changes to the activity patterns in the ventral striatum and amygdala have been observed in humans during contextual fear conditioning (Pohlack et al., 2012), and optogenetic manipulations of neuronal circuits within the amygdala have revealed a bidirectional control of anxiety-related behaviors by these circuits (Tye et al., 2011). Furthermore, increased amygdala activation has been observed in physically healthy humans exhibiting anxious traits (Stein et al., 2007), and maladaptive coupling of the amygdala with other prefrontal circuits may underlie conditions such as post-traumatic stress disorder (PTSD) (Rabinak et al., 2011) and endophenotypes of social anxiety (Bas-Hoogendam et al., 2020). Importantly, these brain regions and defensive behaviors are highly conserved across species (Hur et al., 2020; LeDoux, 2000). Aberrant hyperactivation of the amygdala occurs in some psychological disorders in humans, such as obsessive-compulsive disorder (Simon et al., 2014), attention-deficit hyperactivity disorder (Van Dessel et al., 2020), depression (Peluso et al., 2009), and among PTSD patients (Etkin and Wager, 2007). In general social anxiety disorder, dysregulation of the amygdala creates a bias for threatening social cues, resulting in an increased sensitivity to fearful and angry faces (Labuschagne et al., 2010). Together, our results suggest that AHA^PV^ neurons may recruit downstream circuits to modulate defensive behaviors.

In summary, our work identifies fast-spiking glutamatergic AHA^PV^ neurons as novel regulators of defensive behaviors. Here, we show for the first time that AHA^PV^ neuronal activity is necessary for escape responding. Our findings may help to understand the etiology of some psychophysiological disorders such as aggression, uncontrolled anger, or fear.

## ACKNOWLEDGMENTS

The authors acknowledge with gratitude C. Lupica and D. Wilson for discussions and comments on the manuscript, A. Kisner and the NIDA IRP Histology Core for assistance with radioactive *in situ* hybridizations, S. Lam for assistance with PET scans, and A. Hayden for technical assistance. pAAV-Ef1a-DIO-iC++-EYFP was a gift from K. Deisseroth. Components of the schematic in Figure 1B were created with BioRender. Permission to publish miniscope drawing was granted by Doric Lenses Inc. This work was supported by the National Institute on Drug Abuse Intramural Research Program (NIDA IRP) (ZIADA000595 and ZIADA000069), U.S. National Institutes of Health (NIH).

## AUTHOR CONTRIBUTIONS

Conceptualization B.T.L., M.S.A., A.G., M.M., and Y.A.; Data Curation B.T.L., M.S.A., J.B., A.J., and S.S.; Formal Analysis B.T.L., M.S.A., J.B., and A.J.; Writing – Original Draft B.T.L., M.S.A., and Y.A.; Writing – Review & Editing B.T.L., M.S.A., J.B., A.J., S.S., A.G., M.M., and Y.A.

## REFERENCES

Aponte, Y., Lien, C.-C., Reisinger, E., and Jonas, P. (2006). Hyperpolarization-activated cation channels in fast-spiking interneurons of rat hippocampus. The Journal of Physiology 574, 229–243. https://doi.org/10.1113/jphysiol.2005.104042.

Atasoy, D., Aponte, Y., Su, H.H., and Sternson, S.M. (2008). A FLEX Switch Targets Channelrhodopsin-2 to Multiple Cell Types for Imaging and Long-Range Circuit Mapping. The Journal of Neuroscience 28, 7025. 10.1523/JNEUROSCI.1954-08.2008.

Bang, J.Y., Zhao, J., Rahman, M., St-Cyr, S., McGowan, P.O., and Kim, J.C. (2022). Hippocampus-Anterior Hypothalamic Circuit Modulates Stress-Induced Endocrine and Behavioral Response. Front Neural Circuits 16, 894722. 10.3389/fncir.2022.894722.

Bartholome, O., de la Brassinne Bonardeaux, O., Neirinckx, V., and Rogister, B. (2020). A Composite Sketch of Fast-Spiking Parvalbumin-Positive Neurons. Cerebral Cortex Communications 1, tgaa026. 10.1093/texcom/tgaa026.

Bas-Hoogendam, J.M., van Steenbergen, H., van der Wee, N.J.A., and Westenberg, P.M. (2020). Amygdala hyperreactivity to faces conditioned with a social-evaluative meaning– a multiplex, multigenerational fMRI study on social anxiety endophenotypes. NeuroImage: Clinical 26, 102247. https://doi.org/10.1016/j.nicl.2020.102247.

Boehm, M.A., Bonaventura, J., Gomez, J.L., Solís, O., Stein, E.A., Bradberry, C.W., and Michaelides, M. (2021). Translational PET applications for brain circuit mapping with transgenic neuromodulation tools. Pharmacology Biochemistry and Behavior 204, 173147. https://doi.org/10.1016/j.pbb.2021.173147.

Boudaba, C., Szabo, K., and Tasker, J.G. (1996). Physiological mapping of local inhibitory inputs to the hypothalamic paraventricular nucleus. J Neurosci 16, 7151–7160.

Canteras, N.S., Chiavegatto, S., Ribeiro do Valle, L.E., and Swanson, L.W. (1997). Severe Reduction of Rat Defensive Behavior to a Predator by Discrete Hypothalamic Chemical Lesions. Brain Research Bulletin 44, 297–305. https://doi.org/10.1016/S0361-9230(97)00141-X.

Canteras, N.S., Ribeiro-Barbosa, É.R., and Comoli, E. (2001). Tracing from the dorsal premammillary nucleus prosencephalic systems involved in the organization of innate fear responses. Neuroscience & Biobehavioral Reviews 25, 661–668. https://doi.org/10.1016/S0149-7634(01)00048-3.

Carvalho, V.M.d.A., Nakahara, T.S., Souza, M.A.d.A., Cardozo, L.M., Trintinalia, G.Z., Pissinato, L.G., Venancio, J.O., Stowers, L., and Papes, F. (2020). Representation of Olfactory Information in Organized Active Neural Ensembles in the Hypothalamus. Cell Reports 32, 108061. https://doi.org/10.1016/j.celrep.2020.108061.

Coimbra, N.C., Leão-Borges, P.C., and Brandão, M. (1989). GABAergic fibers from substantia nigra pars reticulata modulate escape behavior induced by midbrain central gray stimulation. Brazilian journal of medical and biological research = Revista brasileira de pesquisas medicas e biologicas 22 *1*, 111–114.

Comoli, E., Ribeiro-Barbosa, É.R., and Canteras, N.S. (2000). Afferent connections of the dorsal premammillary nucleus. Journal of Comparative Neurology 423, 83–98. https://doi.org/10.1002/10969861(20000717)423:1<83::AID-CNE7>3.0.CO;2-3.

Corcoran, K.A., Donnan, M.D., Tronson, N.C., Guzmán, Y.F., Gao, C., Jovasevic, V., Guedea, A.L., and Radulovic, J. (2011). NMDA Receptors in Retrosplenial Cortex Are Necessary for Retrieval of Recent and Remote Context Fear Memory. The Journal of Neuroscience 31, 11655. 10.1523/JNEUROSCI.2107-11.2011.

de Oliveira, L., Hoffmann, A., and Menescal-de-Oliveira, L. (1997). Participation of the Medial and Anterior Hypothalamus in the Modulation of Tonic Immobility in Guinea Pigs. Physiology & Behavior 62, 1171–1178. https://doi.org/10.1016/S0031-9384(97)00351-X.

DeFelipe, J., Lopez-Cruz, P.L., Benavides-Piccione, R., Bielza, C., Larranaga, P., Anderson, S., Burkhalter, A., Cauli, B., Fairen, A., Feldmeyer, D., et al. (2013). New insights into the classification and nomenclature of cortical GABAergic interneurons. Nature reviews. Neuroscience 14, 202–216. 10.1038/nrn3444.

Du, J., Zhang, L., Weiser, M., Rudy, B., and McBain, C.J. (1996). Developmental expression and functional characterization of the potassium-channel subunit Kv3.1b in parvalbumin-containing interneurons of the rat hippocampus. The Journal of Neuroscience 16, 506. 10.1523/JNEUROSCI.16-02-00506.1996.

Esteban Masferrer, M., Silva, B.A., Nomoto, K., Lima, S.Q., and Gross, C.T. (2020). Differential encoding of predator fear in the ventromedial hypothalamus and periaqueductal grey. J Neurosci. 10.1523/JNEUROSCI.0761-18.2020.

Etkin, A., and Wager, T.D. (2007). Functional neuroimaging of anxiety: a meta-analysis of emotional processing in PTSD, social anxiety disorder, and specific phobia. Am J Psychiatry 164, 1476–1488. 10.1176/appi.ajp.2007.07030504.

Falconi-Sobrinho, L.L., and Coimbra, N.C. (2018). The Nitric Oxide Donor SIN-1-Produced Panic-Like Behaviour And Fear-Induced Antinociception Are Modulated By NMDA Receptors In The Anterior Hypothalamus. Journal of Psychopharmacology 32, 711–722. 10.1177/0269881118769061.

Ferris, C.F., Melloni Jr, R.H., Koppel, G., Perry, K.W., Fuller, R.W., and Delville, Y. (1997). Vasopressin/Serotonin Interactions in the Anterior Hypothalamus Control Aggressive Behavior in Golden Hamsters. The Journal of Neuroscience 17, 4331. 10.1523/JNEUROSCI.17-11-04331.1997.

Friedrich, J., Zhou, P., and Paninski, L. (2017). Fast online deconvolution of calcium imaging data. PLOS Computational Biology 13, e1005423. 10.1371/journal.pcbi.1005423.

Fuchs, S.A.G., Edinger, H.M., and Siegel, A. (1985). The role of the anterior hypothalamus in affective defense behavior elicited from the ventromedial hypothalamus of the cat. Brain Research 330, 93–107. https://doi.org/10.1016/0006-8993(85)90010-1.

Hu, H., Gan, J., and Jonas, P. (2014). Interneurons. Fast-spiking, parvalbumin(+) GABAergic interneurons: from cellular design to microcircuit function. Science 345, 1255263. 10.1126/science.1255263.

Hudson, A.E. (2018). Chapter Twelve -Genetic Reporters of Neuronal Activity: c-Fos and G-CaMP6. In Methods in Enzymology, R.G. Eckenhoff, and I.J. Dmochowski, eds. (Academic Press), pp. 197–220. https://doi.org/10.1016/bs.mie.2018.01.023.

Hur, J., Smith, J.F., DeYoung, K.A., Anderson, A.S., Kuang, J., Kim, H.C., Tillman, R.M., Kuhn, M., Fox, A.S., and Shackman, A.J. (2020). Anxiety and the Neurobiology of Temporally Uncertain Threat Anticipation. The Journal of Neuroscience 40, 7949. 10.1523/JNEUROSCI.0704-20.2020.

Kim, D.J., Lee, A.S., Yttredahl, A.A., Gómez-Rodríguez, R., and Anderson, B.J. (2017). Repeated threat (without direct harm) alters metabolic capacity in select regions that drive defensive behavior. Neuroscience 353, 106–118. https://doi.org/10.1016/j.neuroscience.2017.04.012.

Kisner, A., Slocomb, J.E., Sarsfield, S., Zuccoli, M.L., Siemian, J., Gupta, J.F., Kumar, A., and Aponte, Y. (2018). Electrophysiological properties and projections of lateral hypothalamic parvalbumin positive neurons. PLOS ONE 13, e0198991. 10.1371/journal.pone.0198991.

Kwapis, J.L., Jarome, T.J., Lee, J.L., and Helmstetter, F.J. (2015). The retrosplenial cortex is involved in the formation of memory for context and trace fear conditioning. Neurobiology of Learning and Memory 123, 110–116. https://doi.org/10.1016/j.nlm.2015.06.007.

Labuschagne, I., Phan, K.L., Wood, A., Angstadt, M., Chua, P., Heinrichs, M., Stout, J.C., and Nathan, P.J. (2010). Oxytocin Attenuates Amygdala Reactivity to Fear in Generalized Social Anxiety Disorder. Neuropsychopharmacology 35, 2403–2413. 10.1038/npp.2010.123.

Laing, B.J., A; Erbaugh, LJ; Park, AS; Wilson, DJ; Aponte, Y. (2022). Regulation of body weight and food intake by AGRP neurons during opioid dependence and abstinence in mice. Front Neural Circuits, 977642. 10.3389/fncir.2022.977642.

LeDoux, J.E. (2000). Emotion Circuits in the Brain. Annual Review of Neuroscience 23, 155–184. 10.1146/annurev.neuro.23.1.155.

Markram, H., Toledo-Rodriguez, M., Wang, Y., Gupta, A., Silberberg, G., and Wu, C. (2004). Interneurons of the neocortical inhibitory system. Nature reviews. Neuroscience 5, 793–807. 10.1038/nrn1519.

Mendes-Gomes, J., Motta, S.C., Passoni Bindi, R., de Oliveira, A.R., Ullah, F., Baldo, M.V.C., Coimbra, N.C., Canteras, N.S., and Blanchard, D.C. (2020). Defensive behaviors and brain regional activation changes in rats confronting a snake. Behavioural Brain Research 381, 112469. https://doi.org/10.1016/j.bbr.2020.112469.

Michaelides, M., and Hurd, Y.L. (2015). DREAMM: A Biobehavioral Imaging Methodology for Dynamic In Vivo Whole-Brain Mapping of Cell Type-Specific Functional Networks. Neuropsychopharmacology 40, 239–240. 10.1038/npp.2014.233.

Mitra, A., Guèvremont, G., and Timofeeva, E. (2016). Stress and Sucrose Intake Modulate Neuronal Activity in the Anterior Hypothalamic Area in Rats. PLOS ONE 11, e0156563. 10.1371/journal.pone.0156563.

Mittag, J., Lyons, D.J., Sällström, J., Vujovic, M., Dudazy-Gralla, S., Warner, A., Wallis, K., Alkemade, A., Nordström, K., Monyer, H., et al. (2013). Thyroid hormone is required for hypothalamic neurons regulating cardiovascular functions. The Journal of Clinical Investigation 123, 509–516. 10.1172/JCI65252.

Motta, S.C., Goto, M., Gouveia, F.V., Baldo, M.V.C., Canteras, N.S., and Swanson, L.W. (2009). Dissecting the brain’s fear system reveals the hypothalamus is critical for responding in subordinate conspecific intruders. Proceedings of the National Academy of Sciences 106, 4870–4875. 10.1073/pnas.0900939106.

Opland, D., Sutton, A., Woodworth, H., Brown, J., Bugescu, R., Garcia, A., Christensen, L., Rhodes, C., Myers, M., Jr., and Leinninger, G. (2013). Loss of neurotensin receptor-1 disrupts the control of the mesolimbic dopamine system by leptin and promotes hedonic feeding and obesity. Mol Metab 2, 423–434. 10.1016/j.molmet.2013.07.008.

Pacák, J., Točík, Z., and Černý, M. (1969). Synthesis of 2-deoxy-2-fluoro-D-glucose. Journal of the Chemical Society D: Chemical Communications, 77–77. 10.1039/C29690000077.

Peluso, M.A.M., Glahn, D.C., Matsuo, K., Monkul, E.S., Najt, P., Zamarripa, F., Li, J., Lancaster, J.L., Fox, P.T., Gao, J.-H., and Soares, J.C. (2009). Amygdala hyperactivation in untreated depressed individuals. Psychiatry Research: Neuroimaging 173, 158–161. https://doi.org/10.1016/j.pscychresns.2009.03.006.

Pnevmatikakis, E.A., and Giovannucci, A. (2017). NoRMCorre: An online algorithm for piecewise rigid motion correction of calcium imaging data. Journal of Neuroscience Methods 291, 83–94. https://doi.org/10.1016/j.jneumeth.2017.07.031.

Pohlack, S.T., Nees, F., Ruttorf, M., Schad, L.R., and Flor, H. (2012). Activation of the ventral striatum during aversive contextual conditioning in humans. Biological Psychology 91, 74–80. https://doi.org/10.1016/j.biopsycho.2012.04.004.

Qi, J., Zhang, S., Wang, H.L., Barker, D.J., Miranda-Barrientos, J., and Morales, M. (2016). VTA glutamatergic inputs to nucleus accumbens drive aversion by acting on GABAergic interneurons. Nat Neurosci. 10.1038/nn.4281.

Rabinak, C., Angstadt, M., Welsh, R., Kennedy, A., Lyubkin, M., Martis, B., and Phan, K.L. (2011). Altered Amygdala Resting-State Functional Connectivity in Post-Traumatic Stress Disorder. Frontiers in Psychiatry 2, 62.

Root, D.H., Mejias-Aponte, C.A., Zhang, S., Wang, H.L., Hoffman, A.F., Lupica, C.R., and Morales, M. (2014). Single rodent mesohabenular axons release glutamate and GABA. Nat Neurosci 17, 1543–1551. 10.1038/nn.3823.

Rosato-Siri, M.D., Zambello, E., Mutinelli, C., Garbati, N., Benedetti, R., Aldegheri, L., Graziani, F., Virginio, C., Alvaro, G., and Large, C.H. (2015). A Novel Modulator of Kv3 Potassium Channels Regulates the Firing of Parvalbumin-Positive Cortical Interneurons. Journal of Pharmacology and Experimental Therapeutics 354, 251. 10.1124/jpet.115.225748.

Sakurai, K., Zhao, S., Takatoh, J., Rodriguez, E., Lu, J., Leavitt, A.D., Fu, M., Han, B.-X., and Wang, F. (2016). Capturing and Manipulating Activated Neuronal Ensembles with CANE Delineates a Hypothalamic Social-Fear Circuit. Neuron 92, 739–753. https://doi.org/10.1016/j.neuron.2016.10.015.

Schindelin, J., Arganda-Carreras, I., Frise, E., Kaynig, V., Longair, M., Pietzsch, T., Preibisch, S., Rueden, C., Saalfeld, S., Schmid, B., et al. (2012). Fiji: an open-source platform for biological-image analysis. Nat Methods 9, 676–682. 10.1038/nmeth.2019.

Siemian, J.N., Arenivar, M.A., Sarsfield, S., Borja, C.B., Russell, C.N., and Aponte, Y. (2021). Lateral hypothalamic LEPR neurons drive appetitive but not consummatory behaviors. Cell Rep 36, 109615. 10.1016/j.celrep.2021.109615.

Siemian, J.N., Borja, C.B., Sarsfield, S., Kisner, A., and Aponte, Y. (2019). Lateral hypothalamic fast-spiking parvalbumin neurons modulate nociception through connections in the periaqueductal gray area. Scientific Reports 9, 12026. 10.1038/s41598-019-48537-y.

Siemian, J.N., Sarsfield, S., and Aponte, Y. (2020). Glutamatergic fast-spiking parvalbumin neurons in the lateral hypothalamus: Electrophysiological properties to behavior. Physiology & Behavior 221, 112912. https://doi.org/10.1016/j.physbeh.2020.112912.

Simon, D., Adler, N., Kaufmann, C., and Kathmann, N. (2014). Amygdala hyperactivation during symptom provocation in obsessive–compulsive disorder and its modulation by distraction. NeuroImage: Clinical 4, 549–557. https://doi.org/10.1016/j.nicl.2014.03.011.

Sokoloff, L., Reivich, M., Kennedy, C., Rosiers, M.H.D., Patlak, C.S., Pettigrew, K.D., Sakurada, O., and Shinohara, M. (1977). THE [14C]DEOXYGLUCOSE METHOD FOR THE MEASUREMENT OF LOCAL CEREBRAL GLUCOSE UTILIZATION: THEORY, PROCEDURE, AND NORMAL VALUES IN THE CONSCIOUS AND ANESTHETIZED ALBINO RAT1. Journal of Neurochemistry 28, 897–916. https://doi.org/10.1111/j.1471-4159.1977.tb10649.x.

Stein, M.B., Simmons, A.N., Feinstein, J.S., and Paulus, M.P. (2007). Increased Amygdala and Insula Activation During Emotion Processing in Anxiety-Prone Subjects. American Journal of Psychiatry 164, 318–327. 10.1176/ajp.2007.164.2.318.

Thanos, P.K., Robison L Fau - Nestler, E.J., Nestler Ej Fau - Kim, R., Kim R Fau - Michaelides, M., Michaelides M Fau - Lobo, M.-K., Lobo Mk Fau - Volkow, N.D., and Volkow, N.D. (2013a). Mapping brain metabolic connectivity in awake rats with μPET and optogenetic stimulation.

Thanos, P.K., Robison, L., Nestler, E.J., Kim, R., Michaelides, M., Lobo, M.K., and Volkow, N.D. (2013b). Mapping brain metabolic connectivity in awake rats with muPET and optogenetic stimulation. J Neurosci 33, 6343–6349. 10.1523/JNEUROSCI.4997-12.2013.

Tooley, J., Marconi, L., Alipio, J.B., Matikainen-Ankney, B., Georgiou, P., Kravitz, A.V., and Creed, M.C. (2018). Glutamatergic Ventral Pallidal Neurons Modulate Activity of the Habenula–Tegmental Circuitry and Constrain Reward Seeking. Biological Psychiatry 83, 1012–1023. https://doi.org/10.1016/j.biopsych.2018.01.003.

Tye, K.M., Prakash, R., Kim, S.-Y., Fenno, L.E., Grosenick, L., Zarabi, H., Thompson, K.R., Gradinaru, V., Ramakrishnan, C., and Deisseroth, K. (2011). Amygdala circuitry mediating reversible and bidirectional control of anxiety. Nature 471, 358–362. 10.1038/nature09820.

Van Dessel, J., Sonuga-Barke, E., Moerkerke, M., Van der Oord, S., Lemiere, J., Morsink, S., and Danckaerts, M. (2020). The amygdala in adolescents with attention-deficit/hyperactivity disorder: Structural and functional correlates of delay aversion. The World Journal of Biological Psychiatry 21, 673–684. 10.1080/15622975.2019.1585946.

Wallace, M.L., Saunders, A., Huang, K.W., Philson, A.C., Goldman, M., Macosko, E.Z., McCarroll, S.A., and Sabatini, B.L. (2017). Genetically Distinct Parallel Pathways in the Entopeduncular Nucleus for Limbic and Sensorimotor Output of the Basal Ganglia. Neuron 94, 138–152.e135. https://doi.org/10.1016/j.neuron.2017.03.017.

Wang, L., Chen, Irene Z., and Lin, D. (2015). Collateral Pathways from the Ventromedial Hypothalamus Mediate Defensive Behaviors. Neuron 85, 1344–1358. https://doi.org/10.1016/j.neuron.2014.12.025.

Wang, W., Schuette, P.J., Nagai, J., Tobias, B.C., Cuccovia V. Reis, F.M., Ji, S., de Lima, M.A.X., La-Vu, M.Q., Maesta-Pereira, S., Chakerian, M., et al. (2021). Coordination of escape and spatial navigation circuits orchestrates versatile flight from threats. Neuron. https://doi.org/10.1016/j.neuron.2021.03.033.

Yang, C.F., Chiang, M.C., Gray, D.C., Prabhakaran, M., Alvarado, M., Juntti, S.A., Unger, E.K., Wells, J.A., and Shah, N.M. (2013). Sexually dimorphic neurons in the ventromedial hypothalamus govern mating in both sexes and aggression in males. Cell 153, 896–909. 10.1016/j.cell.2013.04.017.

Yilmaz, M., and Meister, M. (2013). Rapid Innate Defensive Responses of Mice to Looming Visual Stimuli. Current Biology 23, 2011–2015. https://doi.org/10.1016/j.cub.2013.08.015.

Zhang, Y. R., Márton; Bushey, Daniel; Zheng, Jihong; Reep, Daniel; Liang, Yajie; Broussard, Gerard Joey; Tsang, Arthur; Tsegaye, Getahun; Patel, Ronak; Narayan, Sujatha; Lim, Jing-Xuan; Zhang, Rongwei; Ahrens, Misha B.; Turner, Glenn C.; Wang, Samuel S.-H.; Svoboda, Karel; Korff, Wyatt; Schreiter, Eric R.; Hasseman, Jeremy P.; Kolb, Ilya;, Looger, Loren L. (2020). jGCaMP8 Fast Genetically Encoded Calcium Indicators. Janelia Research Campus.

Zhou, P., Resendez, S.L., Rodriguez-Romaguera, J., Jimenez, J.C., Neufeld, S.Q., Giovannucci, A., Friedrich, J., Pnevmatikakis, E.A., Stuber, G.D., Hen, R., et al. (2018). Efficient and accurate extraction of in vivo calcium signals from microendoscopic video data. Elife 7. 10.7554/eLife.28728.

