## Supplementary Table 1 for "Anterior hypothalamic parvalbumin neurons are glutamatergic and promote escape behavior"

**Supplementary Table 1. Machine learning model evaluations.**

| Marker-Stage-Model | Sensitivity | Specificity | Accuracy (%) |
|---|---|---|---|
| PV-B-QSVM | 0.846154 | 0.769231 | 75 |
| PV-TE-QSVM | 0.615385 | 1 | 75 |
| PV-A-QSVM | 0.769231 | 0.948718 | 75 |
| PV-E-QSVM | 0.769231 | 0.948718 | 75 |
| PV-B-CosKNN | 0.846154 | 0.692308 | 67.3 |
| PV-TE-CosKNN | 0.692308 | 0.948718 | 67.3 |
| PV-A-CosKNN | 0.769231 | 0.974359 | 67.3 |
| PV-E-CosKNN | 0.384615 | 0.948718 | 67.3 |
| PV-B-CubKNN | 0.692308 | 0.153846 | 23.1 |
| PV-TE-CubKNN | 0.230769 | 0.846154 | 23.1 |
| PV-A-CubKNN | 0 | 0.974359 | 23.1 |
| PV-E-CubKNN | 0 | 1 | 23.1 |
| VGAT-B-QSVM | 0.554217 | 0.851406 | 56.3 |
| VGAT-TE-QSVM | 0.481928 | 0.883534 | 56.3 |
| VGAT-A-QSVM | 0.638554 | 0.863454 | 56.3 |
| VGAT-E-QSVM | 0.578313 | 0.819277 | 56.3 |
| VGAT-B-CosKNN | 0.903614 | 0.951807 | 85.5 |
| VGAT-TE-CosKNN | 0.795181 | 0.959839 | 85.5 |
| VGAT-A-CosKNN | 0.879518 | 0.955823 | 85.5 |
| VGAT-E-CosKNN | 0.843373 | 0.939759 | 85.5 |
| VGAT-B-CubKNN | 0.39759 | 0.907631 | 46.4 |
| VGAT-TE-CubKNN | 0.686747 | 0.606426 | 46.4 |
| VGAT-A-CubKNN | 0.46988 | 0.787149 | 46.4 |
| VGAT-E-CubKNN | 0.301205 | 0.983936 | 46.4 |

**Supplementary Table 1.** Quadratic Support Vector Machine (QSVM), Cosine K-Nearest Neighbor (CosKNN), and Cubic K-Nearest Neighbor (CubKNN) machine learning models were evaluated for sensitivity, specificity, and accuracy of behavioral stage prediction. Models were trained on 80% of the $AHA^{VGAT}$ and $AHA^{PV}$ neurons that were imaged during the predatory threat assay. Marker: PV, $AHA^{PV}$ neurons; VGAT, $AHA^{VGAT}$ neurons. Stages: B, baseline; TE, threat exposure; A, assessment; E, exploration.
